# Understanding Iron and Oxidative Stress Response in *Escherichia coli* Using Multi-phenotype and Ensemble Models

**DOI:** 10.64898/2026.02.04.703689

**Authors:** Daniel Ajuzie, Seyed A. Arshad, Komal S. Rasaputra, Bert Debusschere, Elebeoba E. May

## Abstract

Developing effective antimicrobial strategies requires a predictive understanding of bacterial responses to multiple stress conditions which often result in multiple phenotypes. A microbe’s survival and proliferation depend on its ability to manage concurrent, dynamically varying stressors within its microenvironment. However, time-resolved predictive models that capture multi-phenotype responses are lacking, and single-phenotype models often fail to accurately replicate a microbe’s reaction to mixed stress conditions. In this work, we develop a mechanistic in-silico model of multi-stress response in *Escherichia coli* K12 and use it to characterize phenotype dynamics in iron-limited and hydrogen peroxide containing environments. Specifically, we replicate the iron and oxidative stress response networks in *E. coli* using a system of ordinary differential equations and applied a multi-phenotype parameterization scheme that leverages multi-measure empirical data, augmented metric-based sensitivity analysis, sequential parameter estimation, and ensemble modeling. Our approach resulted in robust models with a 93% accuracy when compared to experimental datasets across 20 stress-response categories, outperforming traditional single-phenotype approaches (80-87%). Analysis of posterior parameter distributions revealed that multi-phenotype optimization eliminates heavy-tailed distributions characteristic of poorly constrained fits and shifts parameter posteriors from boundary-concentrated to centrally localized forms, indicating improved identifiability. Simulation outcomes confirmed key features of *E. coli*’s iron metabolism, showing that moderate peroxide stress in an iron-rich environment creates significant adaptation challenges, leading to a bacteriostatic phenotype. The model provides insights into biochemical mechanisms important to *E. coli*’s temporal response to varying iron availability, with implications for ecological dynamics and pathogenesis. Our parameterization approach highlights the effectiveness of a combination of optimization methods and ensemble modeling in developing predictive models that are robust across multiple phenotypes. Results demonstrate that data structure, specifically the integration of multiple phenotypes and response outputs, proves to be as critical if not more critical than data volume for achieving well-constrained parameter estimates and robust predictions across experimental conditions.

**Author Summary:** Understanding bacterial stress response is crucial for developing strategies to control bacterial populations, particularly as antibiotic resistance poses a growing threat, with "superbugs" potentially triggering a global health crisis. While mathematical models offer powerful tools to study biological systems, many struggle to predict cellular behavior across multiple phenotypes due to the complexity of responses. Iron metabolism is vital for bacterial survival, particularly under oxidative stress, leading to various bacterial growth dynamics. This work uses mathematical modeling to explore how *E. coli* manages multiple stressors, focusing on iron metabolism and oxidative stress. By applying a novel combination of optimization and ensemble modeling methods, we improved model accuracy by nearly 16%, enabling predictions of *E. coli’s* varied response to single, dual, and dynamic stress environments. Our approach offers a valuable tool for understanding and combating bacterial persistence, with future studies able to expand its use to determine how bacterial communities respond to multiple stressors.

## 1. Introduction

Iron is crucial for bacterial metabolism, serving as a cofactor for numerous essential proteins and enzymes. The bio-availability of iron is limited and tightly controlled (1). Ferrous iron (Fe^2+^) is scarce, while ferric iron (Fe^3+^) is largely insoluble and further sequestered by host proteins such as lactoferrin (2),(3),(4). Gram-negative bacteria rely on two iron uptake strategies: permease-mediated import of Fe^2+^ and high-affinity systems – including siderophores and outer-membrane receptors – to capture and reduce Fe^3+^ (3),(4). With a dual and paradoxical role in metabolism, Fe^2+^ is essential as a metabolic cofactor but also fuels Fenton chemistry, generating hydroxyl radicals that damage DNA, proteins, and lipids. These radicals, along with other reactive oxygen species (ROS), can damage several cellular biomolecules and lead to cell death if not maintained at acceptable levels (5). To protect against ROS, bacteria upregulate production of enzymes such as superoxide dismutases (SODs), catalases, and peroxidases (2,6). In mathematical terms, bacteria must balance iron and oxidative fluxes to simultaneously maximize iron uptake to drive metabolic processes while minimizing reactive oxygen species (ROS) production to below toxic thresholds. Understanding how bacteria achieve this balance, which governs bacterial physiology and pathogenicity (1,2,7), will help define new methods for controlling and disrupting bacterial proliferation.

Mechanistic modeling is a powerful tool for deciphering how bacteria manage the complex balance between oxidative stress and iron homeostasis. However, models that explore bacterial responses to multiple stressors, particularly those encountered in dynamic environments (e.g., phagocytic compartments of antigen presenting cells or competitive microbial communities) are few. Developing such models would provide a quantitative understanding of the biochemical interactions that allow bacteria to adapt to fluctuating stressors in their microenvironment, enabling more accurate environment to genotype to phenotype (E ➔G ➔P) predictions.

Several mathematical models addressing iron homeostasis, oxidative stress response, or both have been reported (5,8–18, 19). Semsey et al., developed a kinetic model of iron homeostasis in *E. coli.* This model described bacterial response to iron availability using several uptake and usage pools controlled by the ferric uptake regulator protein (11). More recently, a large metabolic network of *E. coli’s* ROS-based damage and cellular repair processes for metalloproteins was reconstructed by Yang et. al. (5) and was used to explain genome-scale genotype-phenotype relationships associated with stress response. Kinetic reconstructions of the hydrogen peroxide stress network, such as the empirically coupled *E. coli* model by Imlay et al. (15), successfully captured temporal changes in hydrogen peroxide flux. Related models and extensions have been developed, providing insights into specific aspects of the peroxide stress network (5,8–10). These models offer valuable insights into iron metabolism and oxidative stress; however they focus on isolated aspects of the iron and oxidative stress response, which in reality is a co-coordinated process in nearly all bacteria. For example, Semsey’s model thoroughly accounts for extracellular iron levels, uptake, and distribution without incorporating oxidative stress. Imlay’s model focuses on hydrogen peroxide fluxes, with Uhl et al.’s extension adding transcriptional regulation and ROS-associated cell death. However, both models omit iron dynamics and uptake. Yang et al.’s genome-scale model focuses on ROS damage and repair but overlooks other key components of the iron homeostasis network including the effect of kinetic rates on system response (19). To our knowledge, no current model simultaneously addresses extracellular iron dynamics, regulation of iron uptake and usage, and the impact of concurrent and dynamic peroxide induced oxidative stress on *E. coli* metabolism and growth.

### 1.1 Frameworks for Multi-phenotype Model Development

The lack of dynamic, multi-stress models is due in part to the broader challenge of developing robust multi-phenotype models. Kinetic parameters can vary as genetic or environmental conditions change, making it difficult to find a single set of parameters that accurately recapitulates the system across multiple conditions (20,21). To address this challenge, we define an experimental and theoretical framework to quantify and model multi-stress/multi-phenotype systems. First we formalize a definition for phenotype in the context of multi-stress models. A phenotype results from defined environmental conditions, which are produced by a specific combination of stressors (e.g., iron availability and exogenous hydrogen peroxide levels). The environmental condition determines or impacts the physiological and molecular state of the bacteria, resulting in a set of condition-associated, measurable responses or phenotypes (e.g., growth).

For the iron–peroxide stress landscape in *E. coli* (Figure 1A), stress conditions can be divided into four canonical environments bacteria often encounter: (I) low-iron/low-peroxide conditions in niches such as anaerobic colon lumen (22), oxygen-depleted sediments (23), anoxic wastewater bioreactors (24), and sealed fermentation tanks (25); (II) iron-rich but mildly oxidative zones such as the proximal small intestine (26), ferruginous freshwater ponds (27), iron-amended agricultural soils (28), and large-scale bioreactors supplemented with Fe^2+^ (29); (III) conditions where both iron and peroxide are abundant, including inflamed mucosal tissues (30), iron oxide–rich wetland soils (31); and (IV) iron-starved but peroxide-rich environments such as phagolysosome compartments (32), acid-mine drainage sites (33), and oxidant-treated wastewater effluent (34). Each environment often results in quantifiably unique phenotype space characterized by a set of measurable metabolic, genetic, and physiological responses.

**Figure 1:**
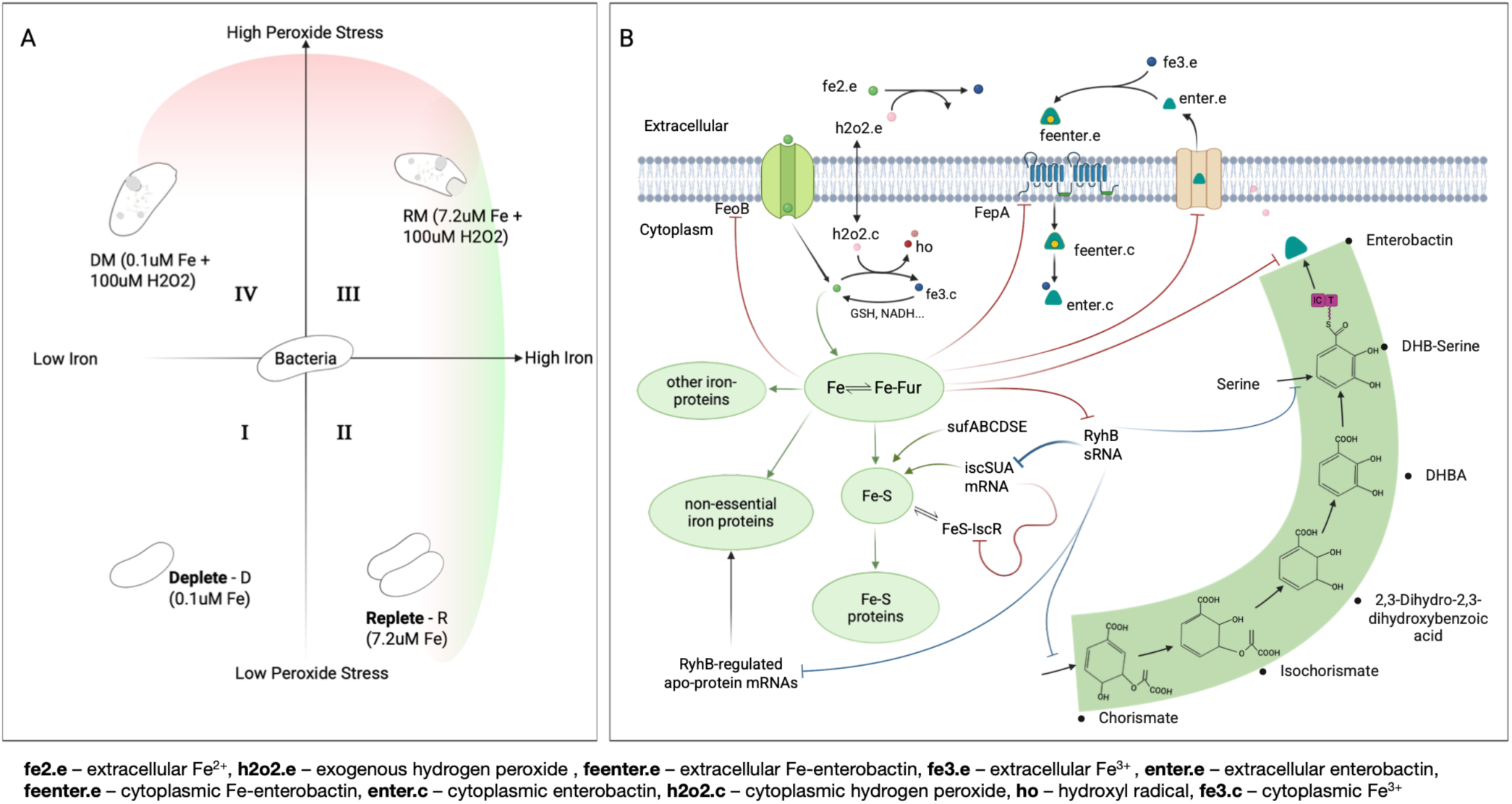
Empirical and Theoretical Frameworks for Regulation of Iron and Peroxide Stress in Escherichia coli. **(A)** Depiction of the two-dimensional multi-stress/multi-phenotype landscape modeled in this work. The two axes represent iron (Fe) availability (left to right) versus exogenous H₂O₂ (reactive oxygen species; ROS) addition (bottom to top), divided into four environment-associated phenotype quadrants: I = low Fe/low ROS, II = high Fe/low ROS, III = high Fe/high ROS, IV = low Fe/high ROS. (B) High-level diagram of E. coli biochemical interactions involved in response to iron/oxidative stress and generation of multi-phenotype outcomes. Species labeled with the subscript “c” are cytoplasmic and those with “e” are extracellular; green arrows denote iron fluxes; red lines indicate transcriptional regulation, and blue lines denote post-transcriptional control.

We developed an experimental model to replicate the four canonical multi-stress conditions and defined their associated multi-phenotype states using a set of experimentally measured outputs, which included: extracellular siderophore concentrations, intracellular and extracellular iron, extracellular hydrogen-peroxide, and biomass accumulation. To build a framework for multi-phenotype theoretical model development, targeting all four iron–peroxide states and drawing on existing stress response models, we reconstructed *E. coli’s* iron/oxidative response network by integrating iron uptake and distribution, H₂O₂ dismutation, gene and protein regulation (e.g., Fur, RyhB, SodA), metabolite dynamics, growth, and extracellular small-molecule exchange (Figure 1B). Our model builds on prior iron and peroxide stress models. The Semsey et al.’s iron regulation model accounts for iron flux using an integrated ferrous and ferric uptake mechanism, with iron distributed into four pools: free- and loosely bound ferrous iron, iron-sulfur cluster and proteins, iron essential and iron non-essential proteins. The Uhl model, extending Imlay’s model, includes peroxide diffusion, catalase and peroxidase detoxification systems, Fenton and redox chemistry of oxidative species, peroxide induced oxidation, and radical (·HO)-induced damage of macromolecules (10). We expand and integrate these two models, using a system of ordinary differential equations (ODEs) to implement a two-compartment replication of *E. coli* multi-phenotype response to dynamic iron/oxidative stress. Ferric iron acquisition is specialized and represented distinctly from ferrous iron uptake. The model includes the *E. coli* enterobactin biosynthesis pathway as it enables high-affinity ferric iron scavenging under iron limitation and has been implicated in mitigating peroxide-induced oxidative stress (35). The model accounts for peroxide influence on iron homeostasis and connects biomolecular stress response to cellular and physiological outcomes such as growth/biomass accumulation.

### 1.2 Parameterization and Use of Multi-phenotype Models

Non-traditional parameterization approaches have been used to define and tune multi-response models, addressing another critical challenge in multi-phenotype modeling. For instance, a *Saccharomyces cerevisiae* iron metabolism model used multi-tiered optimization to iteratively refine parameters across an evolving system of models (14). Sequential parameter estimation has also been shown to improve model accuracy (36) and we previously applied an augmented system of measures to identify influential parameters for single phenotype models (19, 36). However, these methods (9,14,21,36,37) can contain residual sloppiness due to under sampling of the parameter space when optimizing using a single phenotype (38). To address these issues, ensemble modeling (21,39,40), Monte Carlo sampling (37,41,42) and deep learning (43,44) have emerged as alternative and supplemental strategies. These approaches identify multiple parameter sets that span a range of phenotype states, resulting in an optimization process that generates ensembles or sets of parameters capable of approximating empirical data (9,39,41).

In this work, we constructed an *in silico* model to understand how *E. coli* K12 responds and adapts to iron and oxidative stress as single and dual stressors (19). To accurately predict the multiple growth phenotypes that result, we developed a parameterization workflow (Figure 2) that uses an augmented system of measures to identify influential parameters across phenotypes and generate an ensemble of sequentially optimized parameter sets to reproduce time-varying multi-phenotype data (31). To ensure robust parameter calibration while minimizing the risk of overfitting or under-constraining the model, we systematically explored how different configurations of training data impacted optimization performance and model accuracy. Specifically, we implemented sequential optimization for: i) single-phenotype single-response (SPSR) datasets that contain data corresponding to a single measure (e.g., growth CFU) response under a single stress scenario; ii) single-phenotype multiple-response (SPMR) datasets that contain multiple response measures (e.g., growth, extracellular iron) under a single stress scenario; and iii) multiple-phenotype multiple-response (MPMR) datasets that contain multiple response measures (e.g., growth, extracellular iron) under multiple stress scenarios. Sequential optimization allowed us to assess how the accuracy of calibrated models evolved with sequential rounds of parameter optimization. Categorization of the dataset based on stress-associated phenotype groups and measures helped to evaluate whether inclusion of multiple responses or phenotypes was necessary to generate robust models with higher accuracy, or if individual datasets from single phenotype, single response measures could sufficiently constrain the model. The best performing model generated using a standard optimization framework predicts bacterial growth and associated metabolic response under single stress conditions and dual stress conditions with 80% accuracy whether trained using single or multi-stress datasets. However the average accuracy of top performing models was higher for models trained with multiple phenotypes and multiple measures, with top performing MPMR populations having a 6% and 13% advantage in accuracy over SPMR and SPSR, respective. Our model reproduced empirically observed behavior showing *E. coli* is susceptible to iron-associated stress, with comparable susceptibility to a single stress in iron-sufficient conditions (∼7.2 µM) with moderate repeated peroxide stress (∼100 µM H₂O₂) as to dual stress in iron-deficient conditions (<0.01 µM) with the same level of repeated peroxide stress (∼100 µM H₂O₂). Using the model, we explored the mechanistic drivers of single and dual stress-associated differences in growth phenotypes and found that under iron limitation alone, growth slows as *E. coli* upregulates RyhB and siderophore uptake to scavenge scarce iron. In iron-replete cells experiencing oxidative stress, the response pivots to ROS defense, with the system boosting SodA and favoring the more oxidation-tolerant Suf pathway for iron sulfur (Fe–S) cluster assembly. When both stresses coincide, *E. coli* combines these strategies by maximizing iron import (FeoB, FepA), switching from Isc to Suf, and reinforcing antioxidant defenses to protect its Fe–S enzymes and detoxify peroxide; however, growth remains stalled. Results show iron limitation alone reduces growth; iron limitation with oxidative stress ranges from bacteriostatic to bactericidal, whereas iron-replete conditions with oxidative stress is bacteriostatic.

**Figure 2:**
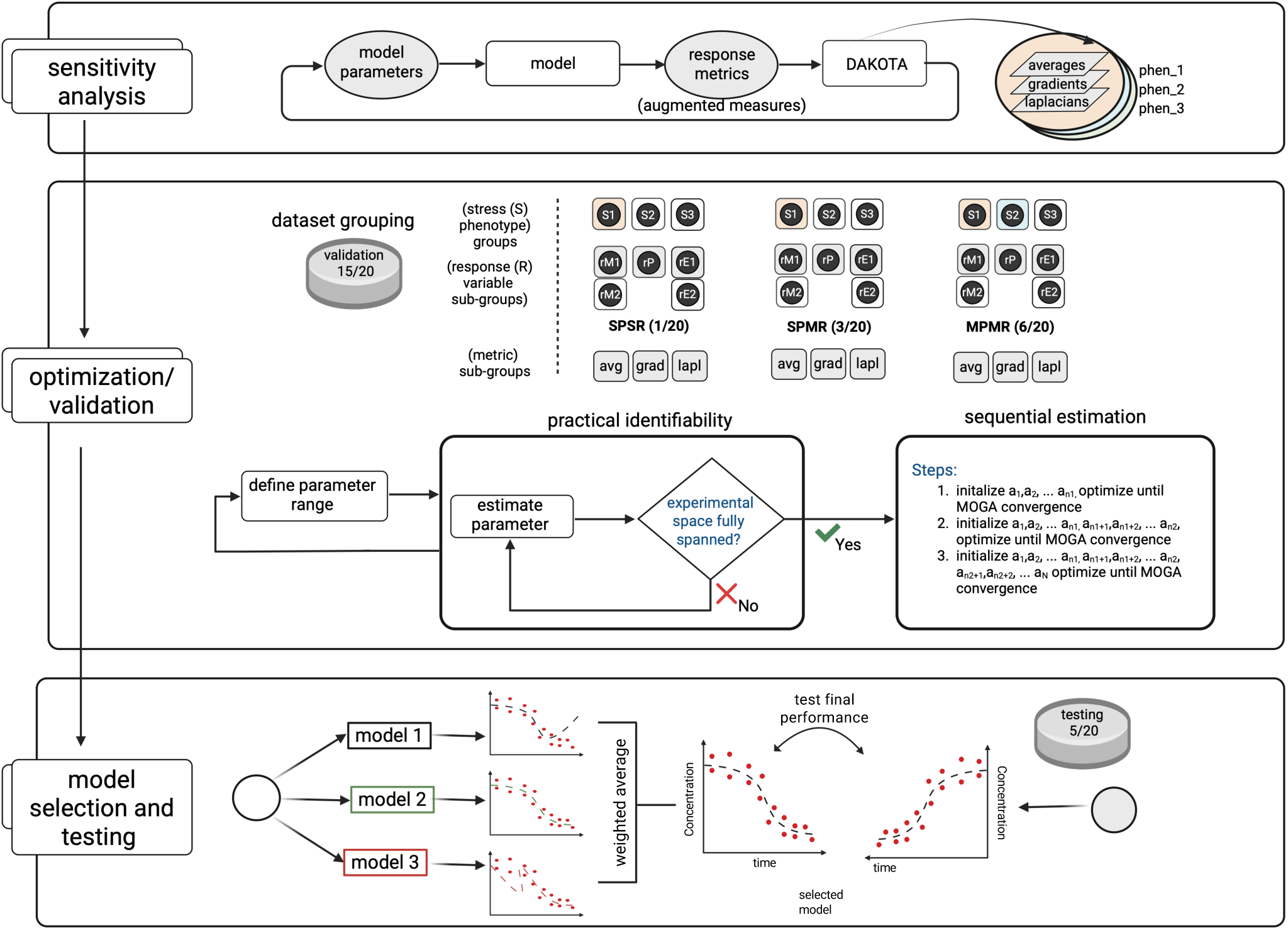
Multi-phenotype Parameterization Workflow. The workflow begins with augmented sensitivity analysis (top panel), which uses the model and DAKOTA, an uncertainty quantification toolkit, to evaluate error metrics using average, gradient, and Laplacians of response measures for each stress phenotype to identify candidate parameters for optimization. During optimization (middle panel) experimental data are partitioned into training/validation and testing sets: three-fourths of the data are used for model training and validation, while one-fourth—drawn from a specific stress phenotype group (S4)—is reserved for testing. Empirical data are grouped by stress phenotypes (S1, S2, S3), each representing distinct environmental conditions that give rise to an associated phenotype, with S4 serving as the test condition or group. Measured response variables are categorized into metabolic (rM; intracellular iron, siderophore), phenotypic (rP; CFU), and environmental (rE; extracellular H_2_O_2_, extracellular iron) response groups. Practical identifiability analysis is then performed to determine feasible parameter bounds. Starting with the most sensitive parameters, model parameters are optimized sequentially across three data and phenotype configurations: SPSR (Single Phenotype Single Response), SPMR (Single Phenotype Multiple Response), and MPMR (Multiple Phenotype Multiple Response). Final model selection (bottom panel) and evaluation are performed using an unweighted and weighted ensemble modeling strategy.

## 2.0 Results and Discussion

### 2.1 Augmented Sensitivity Metrics Identify Robust, Functional Parameter Sets that Drive Steady State and Dynamic System Behaviors

As an alternative to using a single sensitivity metric, we applied multiple, or an augmented system of sensitivity measures to identify and down select a robust set of parameters that capture a majority of the model’s variability (Figure 2, top panel). Measures included nominal averages, gradients (first derivatives) and Laplacians (second derivatives), and were used to identify parameters driving both steady-state and dynamic responses to exogenous stress (19).

Augmented sensitivity analysis (Figure 3; complete results in Figure S1) showed that different sensitivity measures reveal complementary but distinct patterns of parametric influence. These patterns help distinguish parameters that predominantly affect steady-state outcomes from those that drive transient or dynamic behaviors. Some parameters (e.g., volume exchange and iron uptake factors, Figure 3 parameters #1-4,#6: *vole, volc, etvol, binfeob, feobmax*) demonstrate consistent influence across all sensitivity layers, whereas others (e.g., kinetic constants and redox-related rates, Figure 3 parameters #12 - #16: *kffefur, alphafefur, kf, bi, kisc*) exhibit divergence between nominal mean and dynamic measures, particularly between averages and gradient- or Laplacian-based metrics. This divergence suggests that certain parameters are more critical regulators of transient dynamics without exerting strong effects on steady-state outcomes.

**Figure 3:**
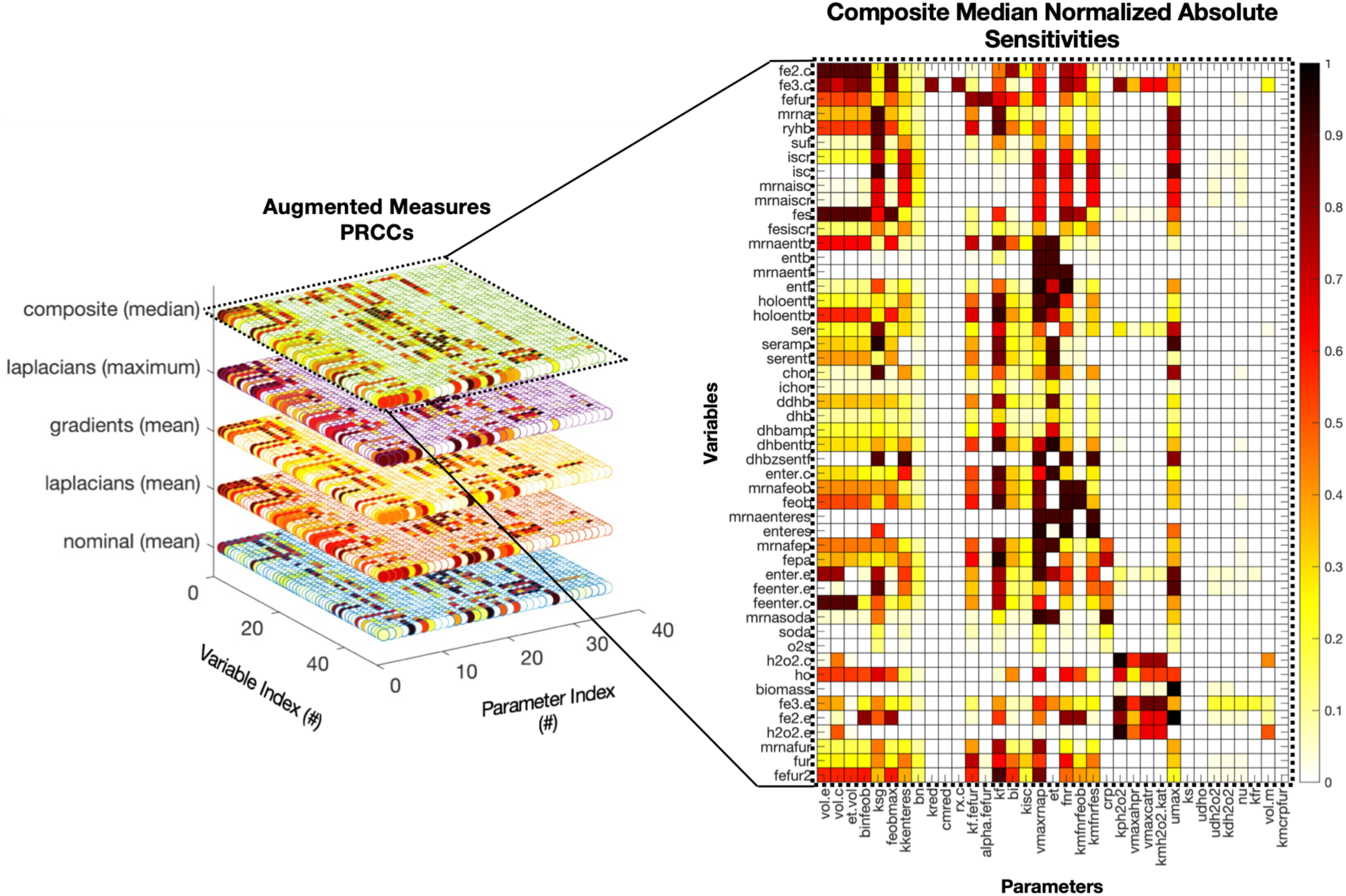
Global Sensitivity Analysis of Iron and Oxidative Stress Model Using Augmented PRCC Measures. Left panel: 3D representation of absolute partial rank correlation coefficients (PRCCs) between model parameters (x-axis) and system responses (y-axis), stratified across four analytical metrics (z-axis): nominal (mean), Laplacians (mean and maximum) and gradients (mean). A fifth layer in z represents the composite sensitivity index computed from median normalized values across the four metrics. Right panel: PRCC heatmap showing monotonic association strength (yellow = low to black = high) between model parameters (columns) and responses (rows). Darker cells (PRCC ≈ 1) pinpoint parameters exerting the greatest influence on iron regulation, oxidative stress, enterobactin biosynthesis, and metabolic intermediates.

In several cases, strong agreement across all sensitivity measures was observed. For instance, intracellular iron levels (variable #1, fe2_c_) showed high sensitivity to volume- and transport-related parameters (parameters #1–4) across all metrics. In contrast, we also observed differences in sensitives. For example, nominal averages resulted in weak associations between fur activation (variables #3, fefur, and #50, fefur2) and iron uptake parameters (#1,#6), whereas gradient and Laplacian measures revealed much stronger correlations (Figure S1). The composite (median) layer aggregates these diverse sensitivity maps, emphasizing parameters that maintain consistent influence across dynamic system and stress conditions. We use the composite metric to facilitate a more robust parameter prioritization approach by reducing dependence on any single sensitivity metric. As shown in the heatmap on the right (flattened view, Figure 3), intracellular iron species (fe2.c, fe3.c, fes) and iron regulatory variables (fefur2, ryhB, iscR) cluster tightly, demonstrating coordinated sensitivity to a shared subset of parameters, primarily parameters governing transport, volume exchange, and iron uptake kinetics. This commonality reflects a structured architecture of regulatory mechanisms governing iron homeostasis, where perturbations to core parameters predictably propagate across related species. In contrast, oxidative stress-related variables, such as h2o2.c, h2o2.e, sodA, and ho, exhibit a dissimilar, scattered sensitivity pattern, suggesting that oxidative stress regulation within the model operates through more modular and condition-specific mechanisms. These observations show the value of using an augmented sensitivity approach to distinguish tightly coupled subsystems, such as iron regulation, from more decentralized modules, such as oxidative stress response. Moreover, results reinforce that while the observed sensitivity patterns are strongly shaped by biochemical network structure, the use of multiple sensitivity measures enables a robust down-selection of influential parameters for optimization.

Following computation of composite sensitivity scores for each stress condition (19, 36), parameters were classified into categories (Table 1) and ranked based on their overall impact on system variables or the percentage of variables the parameter spans: Class I (sensitivity span ≥ 80% of model variables; 14 parameters), Class II (60% ≤ span < 80%; 13 parameters), and Class III (40% ≤ span < 60%; 8 parameters). Analysis of parameter classifications revealed a functional hierarchy: Class I parameters primarily included global response kinetic and regulatory parameters, such as: growth maintenance factor, RNA polymerase transcription rate, Fur dissociation constant, and volumetric scaling factors. Class II parameters were predominantly linked to subsystem regulation, particularly peroxide detoxification and enterobactin biosynthesis pathways. Class III parameters increased granularity and coverage within specific subsystems but contributed less to global variables. Categorization helps differentiate parameters with the greatest composite influence globally versus those that impact subsystems of interest. While not the focus of this work, such insights can be used to reduce complexity of the model prior to subsequent model calibration steps, enabling improvements to both model efficiency and parameter robustness.

**Table 1:**
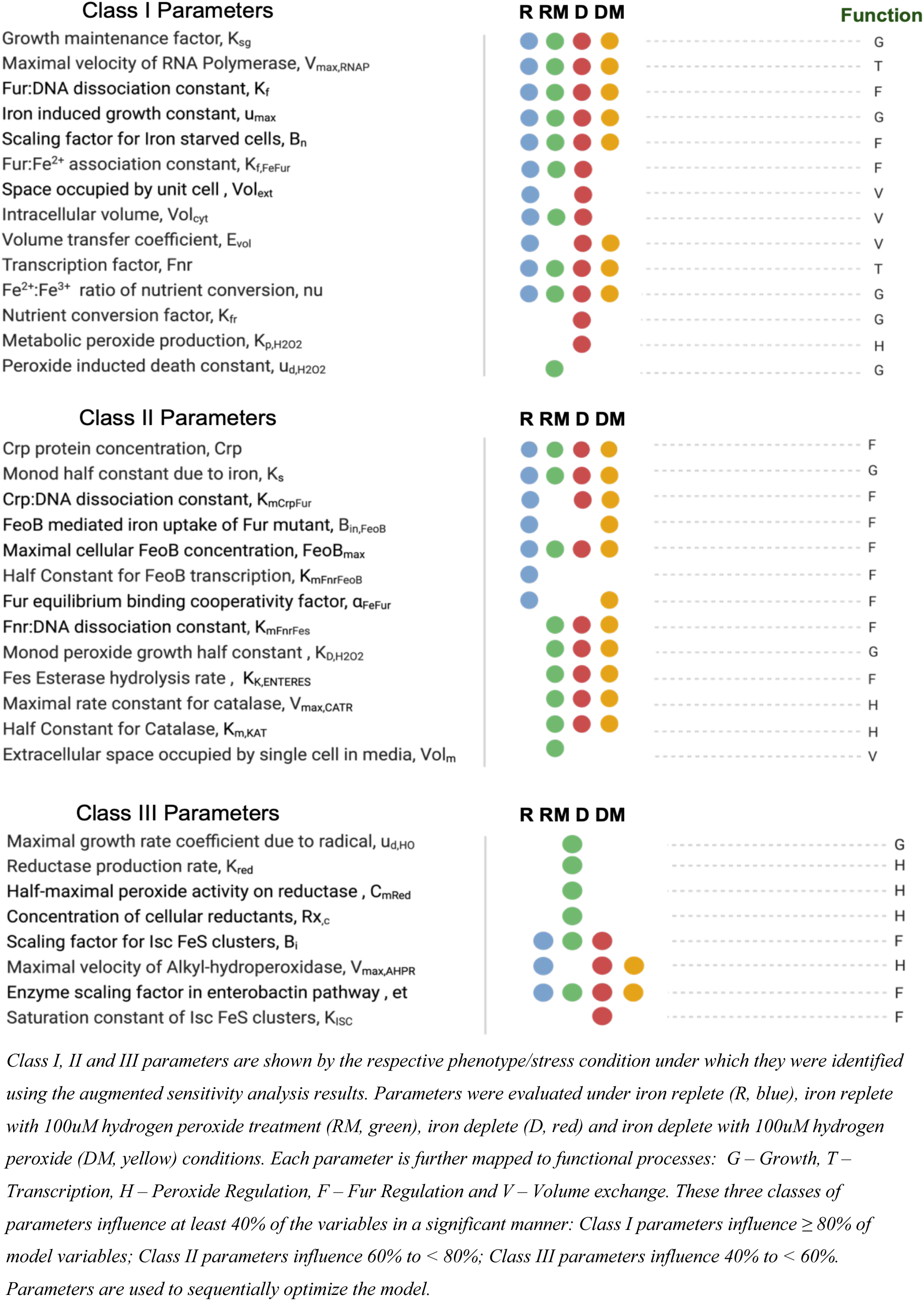
Optimizable Parameters by Classes.

### 2.2 Increasing Dataset Dimensionality Improves the Effectiveness of Sequential Optimization

We used sequential optimization to carry out parameter calibration in stages, first optimizing the most influential set of parameters i.e. Class I parameters, then progressively including Class II and Class III parameters to refine the model further (Figure 2, middle panel). For single phenotype single response datasets (SPSR), optimization produced a single parameter set per phenotype subgroup across 45 possible datasets. In contrast, due to the multi-measure objective metric used for multi-response (SPMR) and multi-phenotype (MPMR) groups, optimization occurred across 36 multidimensional dataset combinations yielding numerous optimal parameter sets per subgroup and hundreds of viable solutions per optimization stage. Figure 4 shows a heatmap of the root mean squared error (log₂(RMSE)) for SPSR, SPMR, and MPMR models, illustrating how sequential parameter refinement improves model fit. While sequential optimization generally improved model fitness, the extent and consistency of improvement varied based on dataset configurations. MPMR consistently decreased model error across all sequential steps and exhibited the most notable improvement in model fitness, with an error reduction of 0.7 log₂(RMSE) for deplete/replete dataset combinations. Although not as robust, SPMR showed improvement and mostly decreased in error for each sequential optimization step. However, SPSR, which used single dimension datasets, had mixed outcomes, with only modest improvements in fitness for some conditions and a decline in fitness for replete datasets. These results show that the impact of sequential optimization is constrained by dataset dimensionality, with the combination of distinct response measures and phenotypes used in MPMR optimization maximizing the effectiveness of multi-stage parameterization.

**Figure 4:**
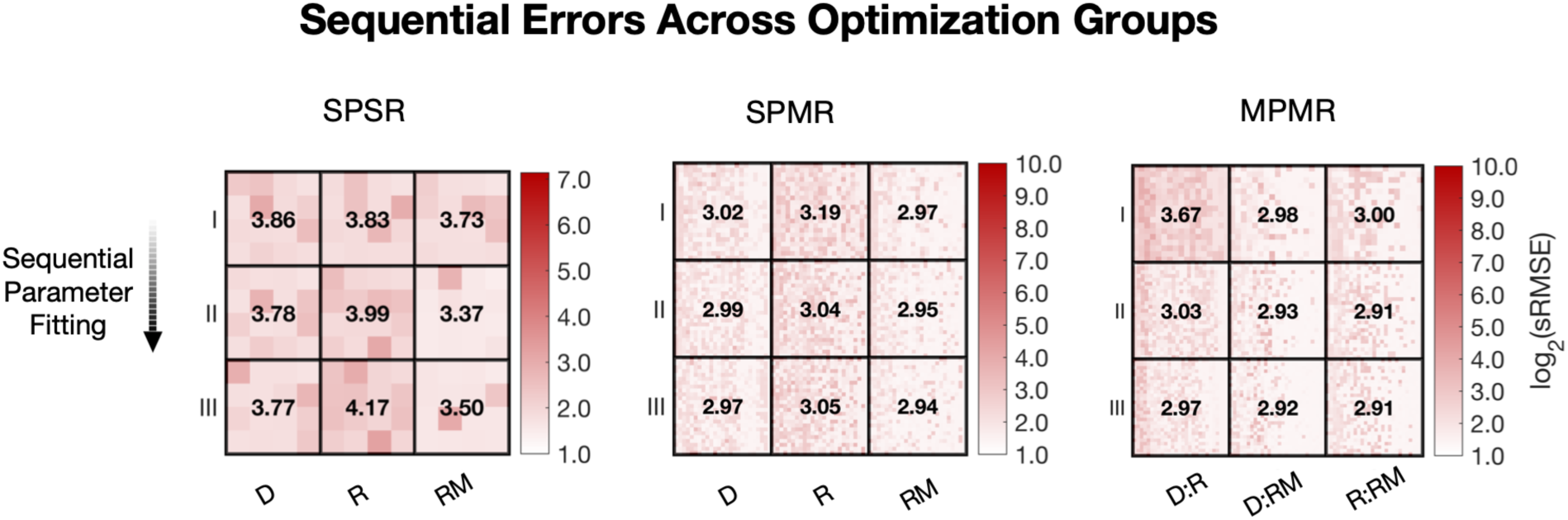
Sequential Estimation Across Optimization Groups. Heatmap showing the relationship between sequential optimization step (rows) and environmental stress conditions (columns) for model prediction error. Cell values represent the mean log₂-transformed sum of range-normalized root mean square error (sRMSE). For MPMR and SPMR, mean sRMSEs shown in cells are computed from the 135 lowest-error values within each sequence-stress combination; for SPSR, all values (15) are used. The pixel mosaic overlay displays individual model errors within each cell. Sequence I contains optimization outcomes from class I parameters alone, Sequence II outcomes are from optimizing class I–II parameters using Sequence I estimates, and Sequence III outcomes are from optimizing class I–III parameters using previous sequence estimates. Stress conditions for SPSR represent individual conditions: (D)eplete, (R)eplete, Replete with 100 µM hydrogen peroxide (RM). SPMR and MPMR use combined conditions (D:R = Deplete and Replete, R:RM = Replete and Replete100, D:RM = Deplete and Replete100). SPSR comprises 3 models × 15 stress–response–metric combinations (45 total), SPMR comprises 136 models × 12 stress–response–metric combinations (1,632 total), and MPMR comprises 138 models × 12 stress–response–metric combinations (1,656 total), each evaluated across 3 stress conditions.

Across all three configurations (SPSR, SPMR, MPMR), sequential optimization from Sequence I (Class I parameters) through Sequence III (Classes I–III) showed distinct, condition-specific improvements in fit. In SPSR, the most significant reduction in error occurred for the iron replete with oxidative stress condition (RM), where median log₂(RMSE) drops from 3.73 in Sequence I to 3.37 and 3.50 in Sequence II and III, respective. This reduction in model error suggests that when using single-phenotype single-response datasets, optimization over broader parameter classes (especially those identified as less sensitive under multi-phenotype screening) can help improve model fit. Additionally, using experimental datasets from stress response phenotypes improves the outcome of optimization.

SPSR optimization using single stress conditions (D and RM) benefited from sequential fitting compared to the non-stressed condition (R). In fact, an increase in error was consistently recorded across optimization sequences unlike with stressed conditions. For SPMR, which introduces additional responses or data dimensions per phenotype, all three conditions benefited substantially from sequential parameter estimation, with iron-replete showing the greatest gains (3.19 to 3.05). For MPMR, which combines phenotype diversity with multiple responses, showed uniform improvements across sequential steps. All three phenotype combinations converged on a minimum log₂(RMSE) by Sequence III, with final errors ranging 2.91–2.97. The highest reduction in model error occurred for the replete:deplete condition, dropping from 3.67 in Sequence I to 2.97 by Sequence III, further showing how multi-phenotype information can help constrain optimization and enable sequential fitting of less sensitive parameters to be more effective. Notably like SPSR, minimum error models for both SPMR and MPMR were obtained for datasets with dynamic stress conditions (replete with medium peroxide stress, RM), with most improvements in fitness occurring between Sequence I and II parameter estimation steps. The relatively flat error landscape in later sequential steps suggests that the model structure, when trained on diverse phenotypes and response measures, becomes well-conditioned and less sensitive to parameter overfitting. These differences in condition-specific outcomes reinforce the use of the augmented sensitivity approach as parameters deemed less globally significant (e.g., in Class III) can still play critical roles in capturing transient or stress-specific behavior, particularly under multiple stress scenarios. The incremental addition of Class III parameters during optimization enables gradual adaptation of the model to condition-specific features, improving model robustness without destabilizing fits in other regions of the parameter space.

### 2.3 Multi-Response Optimization Reduces Error and Increases Robustness of Model Population

We compared MPMR, SPMR, and SPSR optimization strategies by evaluating model errors across phenotype–response combinations and sensitivity metrics (Figure 5). Box plots (Figure 5A-C) display the full distribution of log₂-transformed sRMSE values across all optimized models, while adjacent heatmaps (Figure 5D-F) report median errors from the top 9 lowest error models per condition–response combination (selected to match SPSR cell size: 3 models × 3 metrics), enabling comparison of the best overall performance.

**Figure 5:**
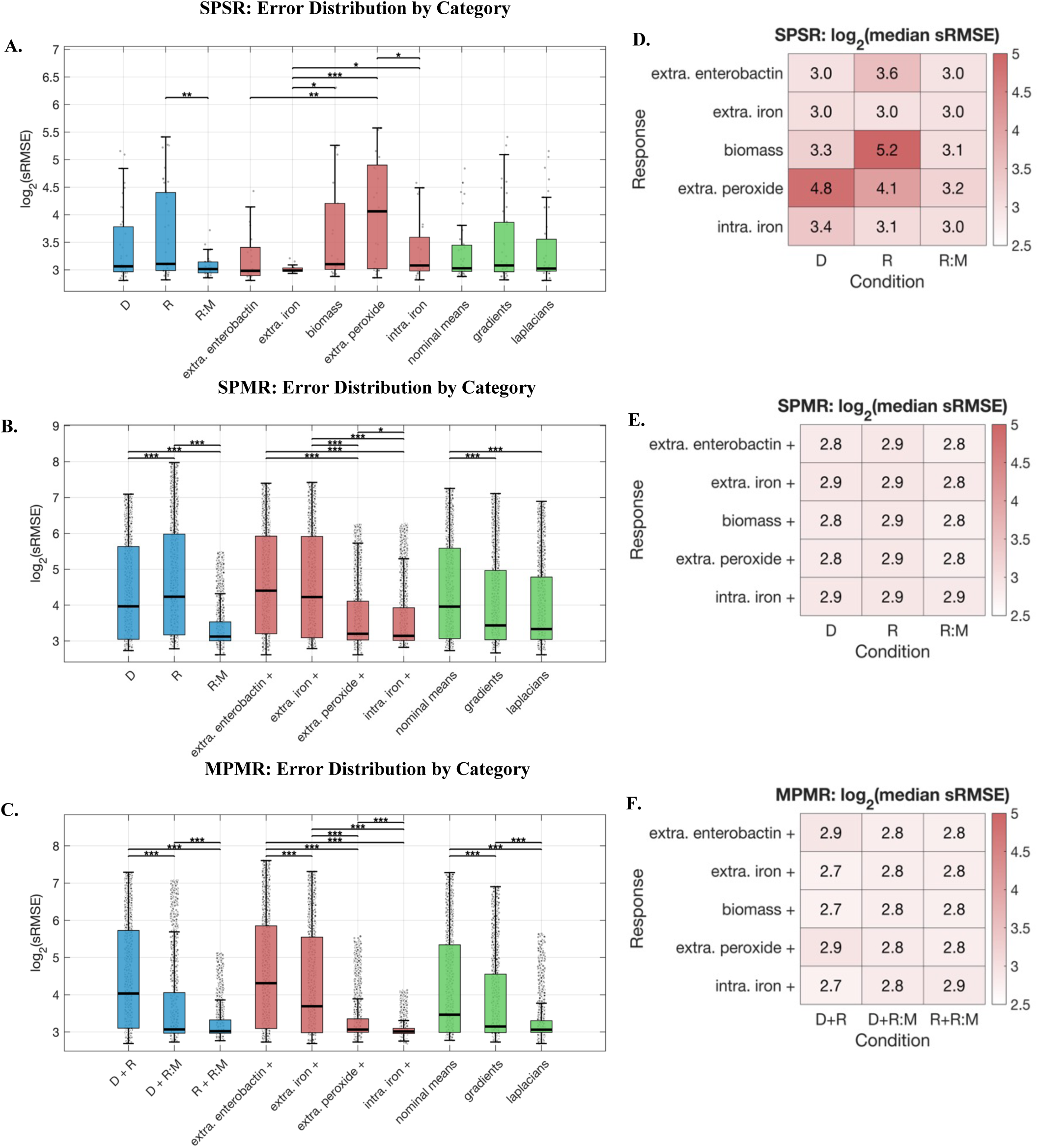
Distribution of model prediction errors by category across optimization modes. (A–C) Box plots of log₂-transformed sum of range-normalized RMSE (sRMSE) grouped by stress condition (blue), response variable (red), and error metric (green) for (A) SPSR, (B) SPMR, and (C) MPMR optimization configurations. Boxes show interquartile range with median; black points denote individual models (jittered). Significance brackets indicate Wilcoxon rank-sum test results (*p < 0.05, **p < 0.01, ***p < 0.001). (D–F) Tables showing log₂-transformed median sRMSE for each response variable (rows) and stress condition (columns) for (D) SPSR, (E) SPMR, and (F) MPMR. Each cell reports the median of the 9 lowest-error models per category, selected to match the SPSR cell size (3 models × 3 metrics). Stress conditions are individual for SPSR/SPMR and paired for MPMR. Response variables include extracellular enterobactin, extracellular iron, extracellular peroxide, intracellular iron, and biomass (biomass always co-optimized in SPMR/MPMR). Responses with a plus sign (e.g extra. iron +) indicate results across all evaluations involving the indicated response. Error metrics include means, gradients, and Laplacians.

Under SPSR configuration, which uses individual phenotype-response pairs, we observed moderate variability in error distributions across condition and response groupings (Figure 5A). Among conditions, R:M (replete with hydrogen peroxide stress) consistently yielded the lowest median errors, while the R (replete) condition produced higher errors. Optimization based on stressed conditions (iron stressed - D and peroxide stress under iron sufficiency - R:M) did not attain statistically different median errors, with both performing better than the unstressed condition - Replete. Response-specific trends (Figure 5D) revealed that extracellular iron and intracellular iron produced relatively lower errors (median ∼3.0), while extracellular peroxide responses exhibited higher errors particularly under iron-deplete conditions (log₂ error of 4.8). Notably, biomass responses under replete conditions showed the highest error among top models in SPSR (log₂ error of 5.2). Among error metrics, Laplacians significantly outperformed both gradients and nominal means (p < 0.001), while gradients showed modest but significant improvement over means (p < 0.05). Critically, even the best-performing SPSR models failed to achieve sub-3.0 error values across most condition–response combinations, suggesting that isolated phenotype-response fitting provides insufficient constraints for robust parameter identification.

In contrast, the SPMR configuration, which captures multiple response outputs for a given phenotype, demonstrated greater variability in log₂(error) across the full model spectrum (Figure 5B) but achieved lower errors for top models (Figure 5E). The replete condition (R) error distribution, while still greater was significantly different from the iron-deplete condition (D) across all models (p < 0.001), with hydrogen peroxide stressed condition producing notably lower error models. This suggests that fitting multiple responses of a system under dynamic stress is a more effective optimization strategy compared to single-stress or single-response (SPSR) fitting, which fails to extract sufficient information from homeostatic conditions. Among response groups, intracellular iron and extracellular peroxide responses showed significantly lower errors compared to extracellular enterobactin and extracellular iron (p < 0.001). Gradients and Laplacian-based optimizations were critical for achieving these results, with both significantly outperforming nominal means (p < 0.001). The heatmap of top-performing models reveals uniformly sub-3.0 median errors across all condition–response combinations (ranging from 2.8 to 2.9), a threshold not achieved by SPSR, demonstrating that optimization using multiple response measures narrows the parameter space in a manner that single-response methods fail to do.

MPMR, which integrates multiple phenotypes and responses in each optimization, provided the clearest separation among conditions, response measures, and metric groups across the full model distribution (Figure 5C). Multi-condition optimization datasets combining iron-deplete and iron-replete phenotypes (D+R) without peroxide stress yielded significantly higher errors compared to both iron-deplete paired with replete plus peroxide stress condition (D+R:M) and iron-replete paired with replete plus peroxide stress (R+R:M) configurations (p < 0.001). This difference indicates that incorporating data from dynamic peroxide stress improves parameter constraint across the model spectrum. Comparing response groups, extracellular enterobactin exhibited significantly higher errors than other responses (p < 0.001), while intracellular iron and extracellular peroxide had the lowest error range. Laplacian-based fits again significantly outperformed gradient and nominal strategies (p < 0.001), reinforcing their suitability for complex, multi-phenotype modeling tasks. The median error for top models (Figure 5F) ranged from 2.7 to 2.9 across all condition–response combinations, representing a modest but consistent improvement over SPMR (2.8–2.9) and a substantial improvement over SPSR (3.0–5.2). This progressive reduction in error demonstrates the benefit of integrating multiple phenotypes and responses to achieve robust, well-constrained models.

Several generalizable principles for calibrating mechanistic models emerge from these results. First, data structure is more critical than data volume. The convergence of top models in SPMR and MPMR to sub-3.0 error values, compared to higher SPSR error values (3.0–5.2) demonstrates that jointly fitting models to multiple phenotypes or response measures helps identify well-constrained parameter regions unattainable through widely-used single measure parameterization methods. Second, system dynamics-based metrics substantially improve calibration quality, with Laplacian and gradient-based optimization consistently outperforming nominal mean approaches; this suggests that experimental designs that perturb the system yields more informative calibration data. Third, the value of perturbation conditions depends partly on data structure. Across all data configurations, fitting using oxidative stress condition produced consistently lower errors with much narrower inter-quartile range compared to other conditions. Deplete condition also performed consistently better that replete condition in both SPSR and SPMR. While stressed conditions outperformed unstressed in SPSR, the SPMR configuration extracted informative constraints even from non-stressed replete conditions by using multiple responses to overcome the limited power of single-measure parameterizations approaches. Fourth, not all response measures are equally effective for model optimization. While some measures (intracellular iron, extracellular peroxide) consistently yielded well-constrained, lower error models, others (enterobactin, biomass) resulted in increased errors. Finally, the significant narrowing of error distributions among best-fit models in multi-phenotype, multi-response (MPMR) configurations indicates that a variation of experimental conditions and response measures are necessary for optimal parameterization and effectively identifying well-constrained parameter regions without overfitting to individual outcome or measures.

### 2.4 Composite Performance Metric Improves Optimal Model Selection

To determine the overall effectiveness of different optimization strategies, we evaluated model coverage alongside error. While error measures the magnitude of deviation between model predictions and experimental observations, its interpretation can vary depending on the response measure and scale, even when normalized. In contrast, coverage provides a scale-independent, statistical measure of model fidelity defined as the proportion of experimental outputs that the model reproduces without statistically significant deviation from observed means (45,46). We assessed coverage across experimentally matched time points and response measures. By applying both metrics – error and coverage – we performed a comprehensive performance analysis across all model optimization strategies. Specifically, we analyzed 4,968 models from 36 dataset groups in MPMR, 4,896 models from 36 groups in SPMR, and 135 models across 45 groups in SPSR. Since SPSR yielded the fewest models, it determined the upper limit for the number of models used in the comparison, and we computed mean, minimum, and maximum performance statistics (Table 2) based on the top 135 models selected under three criteria: error-based (ranked by lowest sRMSE), coverage-based (ranked by highest coverage), and composite (ranked by a balanced score weighting both metrics equally).

**Table 2:** Model Performance Across Optimization Strategies and Metrics.

| Performance Metric | Error-based |  |  | Coverage-based |  |  | Composite |  |  |
| --- | --- | --- | --- | --- | --- | --- | --- | --- | --- |
|  | MPMR | SPMR | SPSR | MPMR | SPMR | SPSR | MPMR | SPMR | SPSR |
| **N=1 (single model no error biasing)** |  |  |  |  |  |  |  |  |  |
| minimum error | 6.45<sup>a</sup> | 6.13<sup>c</sup> | 7.01<sup>a</sup> | 6.54<sup>a</sup> | 6.80<sup>a</sup> | 7.01<sup>a</sup> | 6.54<sup>a</sup> | 6.13<sup>c</sup> | 7.01<sup>a</sup> |
| corresponding coverage | 0.67 | 0.8 | 0.73 | 0.8 | 0.8 | 0.73 | 0.8 | 0.8 | 0.73 |
| mean error | 7.17 | 7.23 | 22.79 | 24.57 | 26.28 | 22.79 | 18.24 | 14.05 | 22.79 |
| mean coverage | 0.65 | 0.68 | 0.6 | 0.79 | 0.8 | 0.6 | 0.79 | 0.8 | 0.6 |
| maximum coverage | 0.80<sup>ac</sup> | 0.80<sup>ac</sup> | 0.80<sup>abc</sup> | 0.93<sup>a</sup> | 0.80<sup>abc</sup> | 0.80<sup>abc</sup> | 0.93<sup>a</sup> | 0.80<sup>abc</sup> | 0.80<sup>abc</sup> |
| corresponding error | 6.54 | 6.13 | 7.9 | 29.79 | 6.8 | 7.9 | 29.79 | 6.13 | 7.9 |
| **N=1 (single model with error biasing)** |  |  |  |  |  |  |  |  |  |
| minimum error | 6.81<sup>c</sup> | 7.10<sup>a</sup> | 7.01<sup>a</sup> | 29.79<sup>a</sup> | 7.10<sup>a</sup> | 7.01<sup>a</sup> | 6.81<sup>c</sup> | 7.10<sup>a</sup> | 7.01<sup>a</sup> |
| corresponding coverage | 0.8 | 0.87 | 0.73 | 0.93 | 0.87 | 0.73 | 0.8 | 0.87 | 0.73 |
| mean error | 7.09 | 7.87 | 7.21 | 29.79 | 9.36 | 9.84 | 7.13 | 8.43 | 8.48 |
| mean coverage | 0.8 | 0.76 | 0.67 | 0.93 | 0.81 | 0.74 | 0.8 | 0.81 | 0.74 |
| maximum coverage | 0.80<sup>c</sup> | 0.87<sup>a</sup> | 0.80<sup>a</sup> | 0.93<sup>a</sup> | 0.87<sup>ab</sup> | 0.80<sup>ac</sup> | 0.87<sup>b</sup> | 0.87<sup>ab</sup> | 0.80<sup>ac</sup> |
| corresponding error | 6.81 | 7.1 | 7.03 | 29.79 | 7.1 | 7.03 | 9.03 | 7.1 | 7.03 |
| **N>1 (ensemble with error biasing)** |  |  |  |  |  |  |  |  |  |
| minimum error | 6.81<sup>c</sup> | 6.88<sup>a</sup> | 6.69<sup>a</sup> | 6.98<sup>c</sup> | 6.97<sup>a</sup> | 7.00<sup>a</sup> | 6.81<sup>c</sup> | 6.90<sup>a</sup> | 6.90<sup>a</sup> |
| corresponding coverage | 0.8 | 0.73 | 0.73 | 0.87 | 0.8 | 0.8 | 0.8 | 0.8 | 0.8 |
| mean error | 6.91 | 6.98 | 6.91 | 25.22 | 10.32 | 9.26 | 6.94 | 7.49 | 7.42 |
| mean coverage | 0.8 | 0.76 | 0.66 | 0.87 | 0.81 | 0.8 | 0.81 | 0.81 | 0.8 |
| maximum coverage | 0.80<sup>c</sup> | 0.80<sup>ac</sup> | 0.80<sup>a</sup> | 0.93<sup>a</sup> | 0.87<sup>abc</sup> | 0.87<sup>c</sup> | 0.87<sup>bc</sup> | 0.87<sup>abc</sup> | 0.87<sup>c</sup> |
| corresponding error | 6.81 | 6.9 | 6.9 | 18.31 | 7.06 | 7.28 | 6.98 | 7.06 | 7.28 |

Performance metrics for three optimization approaches (MPMR, SPMR, SPSR) evaluated under three model selection criteria: Error-based (top 135 models ranked by lowest sum of range-normalized RMSE), Coverage-based (top 135 models ranked by highest coverage proportion), and Composite (top 135 models ranked by composite score = $0.5 \times (1 - \text{normalized error}) + 0.5 \times \text{normalized coverage}$ , where normalization maps values to $[0,1]$ range). Considering single versus ensemble models and use of error biasing, three model categories are compared: $N=1$ without error biasing, $N=1$ with error biasing, and ensemble i.e. $N>1$ with error biasing where $N$ is the number of models in the ensemble. Maximum number of models in ensembles was set to 108 based on piecewise mapping to experimental data. Metrics reported: minimum error (lowest sRMSE among top 135 models), corresponding coverage (highest coverage proportion among minimum error models), mean error and mean coverage (arithmetic means of top 135 model), maximum coverage (highest coverage proportion among top 135 models), and corresponding error (lowest sRMSE among maximum coverage models). Superscript letters indicate stress condition dataset used for optimization: $a$ = Deplete:Replete (or Deplete for SPSR, SPMR), $b$ = Replete:Replete100 (or Replete for SPSR, SPMR), $c$ = Deplete:Replete100 (or Replete100 for SPSR, SPMR). Multiple letters indicate ties across stress groups.

Results from the baseline optimization approach (N=1, no error biasing),, where we evaluate optimized models against the entire validation datasets (N=1), the choice of model selection criterion substantially influenced performance outcomes. When models were selected based on error, SPMR achieved the lowest minimum error (6.13) with corresponding coverage of 0.80, followed by MPMR (6.45, coverage 0.67) and SPSR (7.01, coverage 0.73). SPSR average performance was three times worse, with a higher mean error (22.79) compared to MPMR (7.17) and SPMR (7.23); this reflects the limitations of using single phenotype-response fitting. Mean coverage for error-based selection followed a similar pattern, with SPMR (0.68) and MPMR (0.65) outperforming SPSR (0.60).

Coverage-based selection revealed an inherent trade-off: while MPMR achieved the highest maximum coverage (0.93), this came at the cost of substantially elevated error (29.79), whereas SPMR maintained both high coverage (0.80) and low corresponding error (6.80). The composite selection criterion balanced these trade-offs, with SPMR achieving both the lowest minimum error (6.13) and high coverage (0.80). Introduction of error biasing (N=1, with error biasing) improved coverage across all optimization modes while maintaining competitive error values. MPMR achieved the highest mean coverage (0.80) with a mean error of 7.09, while SPMR attained higher maximum coverage (0.87) with corresponding error of 7.10. Notably, coverage-based selection for MPMR produced a model with exceptional coverage (0.93) but poor error performance (29.79), highlighting that optimizing solely for coverage can yield models that statistically match experimental means while failing to minimize temporally dependent prediction error. SPSR models showed improved mean coverage (0.67–0.74) with error biasing but remained below the coverage levels achieved by multi-response configurations.

Ensemble-based evaluations (N>1, with error biasing) further improved performance metrics and narrowed differences among optimization approaches. Under error-based selection, all three configurations achieved comparable minimum errors (MPMR: 6.81, SPMR: 6.88, SPSR: 6.69), with mean errors converging to similar values (6.91–6.98). This convergence suggests that ensemble approaches can partially compensate for inherent weakness in SPSR by aggregating predictions across multiple models. However, coverage metrics continued to differ across configurations. Even with ensemble methods, MPMR maintained the highest maximum coverage (0.93 under coverage-based selection) with mean coverage of 0.80–0.87, while SPSR had lower mean coverage (0.66–0.80) despite competitive error values. Under composite selection, MPMR achieved the best balance of minimum error (6.81) and maximum coverage (0.87), with SPMR and SPSR showing similar but slightly reduced performance.

These results demonstrate the challenge of balancing error minimization for individual time points and overall data coverage maximization, with the composite selection criterion providing a practical approach for selecting models that balance both objectives. The more standard SPSR approach returns weaker performing models. However, incorporating ensemble methods with error biasing can improve outcomes, yielding competitive SPSR model sets with minimum errors. MPMR and SPMR dataset configurations and optimization strategies cover a broader solution space and increase the likelihood of identifying higher performance models with robust parameter sets able to effectively cover a range of phenotypes.

### 2.6 Validated Multi-phenotype Models Predict Response to Dual Stress Condition

Model validation was performed by selecting a single best-fit model from each configuration (SPSR, SPMR and MPMR) using the composite scoring system that combined error and coverage metrics to determine the top scoring model. Top scoring models do not necessarily have the lowest error or the maximum coverage but are balanced to capture dynamics (error) and overall response (coverage). Using the selected best-fit models we tested each model’s ability to reproduce five key outputs – cell density, intracellular iron, enterobactin, extracellular iron, and peroxide – across three conditions (replete, deplete, replete + peroxide). As depicted in Figure 6, the MPMR model delivered the most reliable predictions, achieving ∼73% coverage for the validation set (R, D, R:M), and accurately tracked both intracellular and extracellular iron levels under single-stress conditions. The SPMR model, with a ∼67% coverage, effectively captured metabolic response measures (iron pools, enterobactin), but showed mixed performance on growth and extracellular peroxide variables. SPSR with a coverage of ∼53% exhibited the greatest discrepancies between prediction and experiment. Notably, all strategies had challenges predicting *E. coli* response under replete plus hydrogen peroxide (RM) stress, resulting in 20% coverage for that condition.

**Figure 6.**
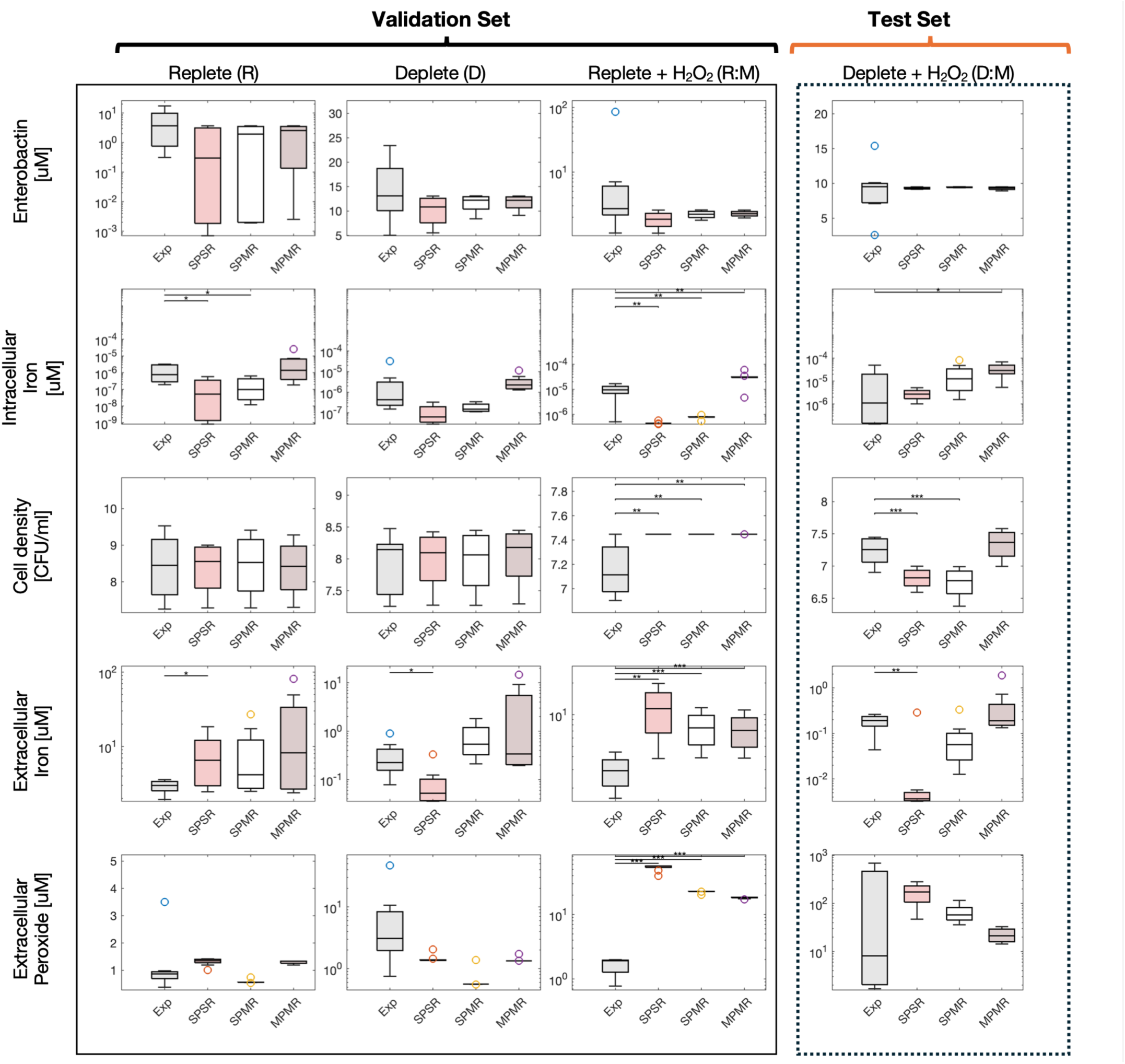
Model Validation Across SPSR, SPMR, and MPMR Configurations Using Experimental Data from E. Coli K12 Grown Under Various Iron and Oxidative Stress Conditions. The boxplots compare model predictions (colored) with experimental data (gray), showing model performance under replete, deplete, and replete with peroxide conditions. Statistical significance is indicated by asterisks, highlighting significant deviations between model predictions and experimental data (* p ≤ 0.05; ** p ≤ 0.01; *** p ≤ 0.001).

Next, we tested the models against a new dual-stress condition (D:M) to determine model robustness and ability to predict *E. coli* behavior in more complex environments. Both MPMR and SPMR models achieved 80% coverage whereas the SPSR model reached only 60% coverage.

Differences in model performance highlights how optimization and training strategy affects the ability to generalize across different phenotypes and responses. MPMR parameterization, integrating multiple phenotypes and responses, performed well across the majority of responses but had a slight challenge predicting intracellular iron under deplete-medium stress conditions. SPMR, which includes multiple responses for each phenotype, produced generalizable models comprable to MPMR in coverage for the test datasets. As expected, the SPSR configuration, relying on single-response data, was less able to generalize as effectively, with a 60% coverage of the test datasets. Use of ensemble models improved SPSR coverage for both the validation and test datasets (ensemble validation is shown in Supplementary S2 Figure).

### 2.7 Ensemble Model Mechanistically Demonstrates How Oxidative Stress Drives Distinct Regulatory Shifts in Iron Homeostasis

To improve prediction of dynamic response and the temporal fit of the model, we used linear programming methods to identify and evaluate an optimal ensemble model from a population of candidate models from each parameterization approach (SPSR, SPMR, and MPMR, Supplementary Figure S2). The MPMR ensemble, composed of three candidate models, effectively captured the overall and dynamic response of *E. coli* under multiple stress phenotypes. Figure 7 shows the optimal MPMR ensemble’s temporal profiles for sixteen key molecular and physiological response variables (the remaining 34 response variables are plotted in Supplementary Figures S3 – S4) across four environmental conditions: iron-deplete (D), iron-replete (R), iron-deplete with repeated 100µM hydrogen peroxide (DM), and iron replete with repeated 100µM hydrogen peroxide (RM). Simulations represent a 12-hour response window corresponding to the length of the experimental study. Notably, the MPMR ensemble captures both qualitative and quantitative trajectories more accurately than SPMR, particularly for Fe–S/IscR, Fur, and oxidative defense subsystems (51),(52),(11). Table 3 summarizes these observed responses across environmental phenotypes.

**Figure 7.**
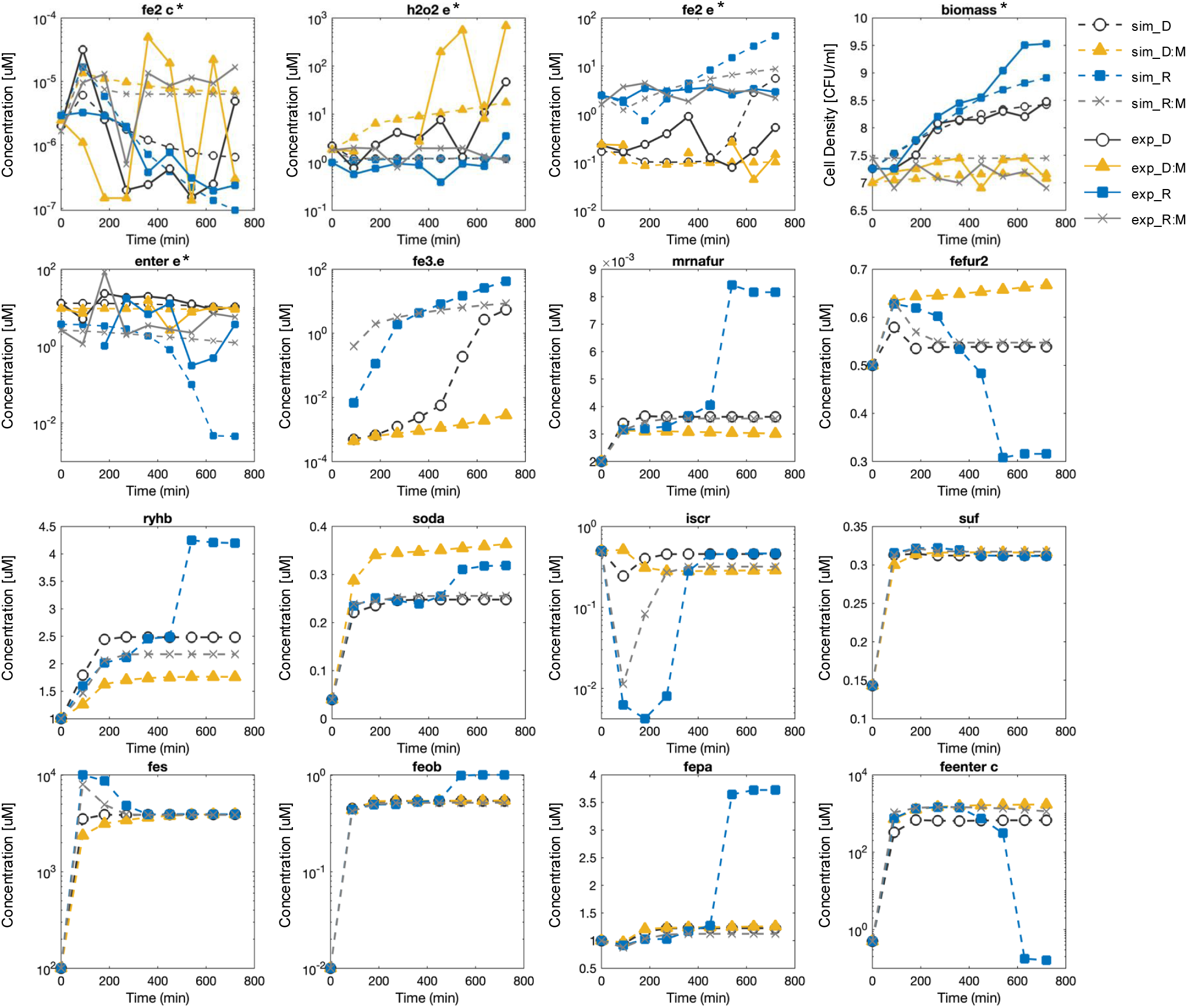
Simulated vs. experimental time courses under four stress phenotypes (D, DM, R, RM). Dotted lines show model-predicted trajectories for 16 variables—extracellular Fe²⁺ (fe2.e), extracellular H₂O₂ (h2o2.e), intracellular Fe²⁺ (fe2.c), biomass, extracellular enterobactin (enter.e), extracellular Fe³⁺ (fe3.e), fur mRNA (furmrna), active Fur protein (fefur2), RyhB sRNA (ryhb), SodA (soda), IscR regulator (iscr), Suf pathway proteins (suf), Fe–S clusters (fes), FeoB transporter (feoB), FepA transporter (fepa), and intracellular Fe–enterobactin (feenter.c). Solid lines overlay experimental measurements for the first five variables marked with asterisks - fe2.e, h2o2.e, fe2.c, biomass, enter.e under iron-deplete (D), iron-deplete + 100 µM H₂O₂ (DM), iron-replete (R), and iron-replete + 100 µM H₂O₂ (RM) conditions.

**Table 3:** Summary of Simulated Molecular and Physiological Responses to Iron and Oxidative Stress in Escherichia coli. Qualitative (and quantitative) descriptors summarize model-predicted responses across four conditions. Iron-replete without stress (R) serves as the physiological baseline; all other conditions are compared to R, with percentage changes shown in parentheses. Tabular data reveals how iron limitation (D), oxidative stress (RM), and their combination (DM) each perturb homeostasis relative to the unstressed, iron-sufficient state. Iron transporters and extracellular enterobactin are normalized per cell (value divided by biomass) to account for population size differences across conditions. ᵃ Per-cell normalized: concentration divided by biomass to account for population size differences. All values are averages (t = 0–720 min). Percentages indicate change relative to R (iron-replete, no stress) baseline.

| Response | Species<br>(Variable Name) | Iron-replete<br>(R) | Iron-deplete<br>(D) | Iron-replete +<br>H<sub>2</sub>O<sub>2</sub> (RM) | Iron-deplete +<br>H<sub>2</sub>O<sub>2</sub> (DM) |
| --- | --- | --- | --- | --- | --- |
| Iron pools | Extracellular<br>Fe<sup>2+</sup> (fe2.e) | Abundant | Scarce | Abundant | Scarce |
|  | Intracellular labile<br>Fe<sup>2+</sup> (fe2.c) | Elevated pool | Reduced pool | Moderately<br>reduced | Severe depletion |
|  | Extracellular<br>Fe<sup>3+</sup> (fe3.e) | Moderate<br>oxidation | Low level | Consumed | Consumed |
| Fur<br>regulation | fur mRNA<br>(mrnafur) | Stable<br>expression | Suppressed<br>expression | Strong<br>suppression | Strong<br>suppression |
|  | Active holo-Fur<br>(fefur2) | Basal activity | Increased activity | Increased activity | Sustained<br>activation |
|  | IscR<br>regulator (iscr) | Repressed | Derepressed | Slightly<br>repressed | Partially<br>derepressed |
|  | Suf pathway (suf) | Stable | Maintained | Stable | Maintained |
|  | Fe-S<br>clusters (fes) | Well<br>maintained | Depleted | Destabilized | Strongly<br>depleted |
| Iron<br>transporters <sup>a</sup> | FeoB/cell<br>(feob) | Basal | Elevated | Strongly<br>induced | Maximally<br>induced |
|  | FepA/cell<br>(fepa) | Basal | Slightly<br>elevated | Strongly<br>induced | Maximally<br>induced |
| Siderophore<br>system | Intracellular<br>Fe-enterobactin<br>(feenter.c) | Low<br>accumulation | Lower | Elevated | Strongly<br>elevated |
|  | Extracellular<br>enterobactin/cell <sup>a</sup><br>(enter.e) | Basal<br>secretion | Strong<br>secretion | Reduced<br>secretion | Maximal<br>secretion |
| Regulatory<br>RNA | RyhB<br>(ryhb) | Induced | Reduced | Suppressed | Strongly<br>suppressed |
| ROS<br>defense | SodA<br>(soda) | Moderate<br>expression | Low<br>expression | Low<br>expression | Induced |
| Physiology | Cell density<br>(biomass) | Steady<br>growth | Slower<br>growth | Growth<br>plateau | Decline/<br>collapse |

We compared simulation outputs with experimental data for the five measured variables. Under iron-sufficient conditions (R, RM), extracellular iron (fe2.e, fe3.e) is readily available for uptake (47) whereas it is scarce under iron-depleted conditions (D, DM). In deplete, oxidative stress compounds iron limitation by shifting speciation from Fe²⁺ toward less bioavailable Fe³⁺, deepening the supply constraint. In replete, where iron is abundant, oxidative stress does not significantly impair bioavailability but instead disrupts downstream iron-dependent processes such as utilization and redox-sensitive regulation via altered speciation and signaling. Model outputs show that oxidative stress transiently elevates reactive Fe²⁺ in labile intracellular pools (fe2.c), while cumulatively depleting iron from catalytic Fe–S cluster pools (fes), which are vulnerable to oxidative degradation.

Hydrogen peroxide dynamics further distinguish iron-sufficient oxidative stress (RM) from dual stress (DM). In DM, peroxide is initially buffered but eventually accumulates, overwhelming cellular defenses and driving biomass attenuation. In RM, peroxide rises sharply but is progressively detoxified, reflecting active stress mitigation; however, biomass remained attenuated as well. Thus, although cells under dual stress (DM) handle oxidative insult less effectively than those under oxidative stress alone (RM), the physiological outcome is similar. Iron-starved cells (D) also exhibit limited capacity to manage internal oxidants, showing elevated peroxide levels at later stages compared to non-stressed cells (R). Labile Fe²⁺ (fe2.c) shows transient oscillations under oxidative stress (RM, DM) but is depleted on average compared to unstressed conditions (R, D), with the most severe depletion under dual stress (DM, −33%). However oxidative stress (RM, DM) increased Fe²⁺ at later timepoints, which is captured by the model. These instabilities underscore the acute sensitivity of the bacteria to combined iron limitation and oxidative challenge.

The observed dynamics suggest that bacteria may perceive oxidative stress in iron-rich environments (RM) as functionally equivalent to iron-starvation despite abundant extracellular Fe, possibly due to a Fenton-related shift toward an increase in bioavailable Fe³⁺. Iron speciation and redox dynamics emerge as key modulators of cellular stress responses, determining the cell’s ability to maintain redox balance and iron homeostasis. This interplay is reflected in the behavior of Fur, the master regulator of iron metabolism. Active holo-Fur (fefur2), which requires Fe²⁺ for DNA binding, shows partial activation under single stress but reaches full activation under DM despite low transcript levels suggesting a compensatory shift toward more reactive Fe. Interestingly, holo-Fur dynamics under RM and D are remarkably similar, potentially explaining shared downstream metabolite profiles between these stress phenotypes. RyhB, a small RNA that normally helps regulate iron by repressing the synthesis of non-essential iron-containing proteins during iron limitation (D), is suppressed in RM despite iron abundance and even more so under DM conditions. In this ensemble model, RyhB levels are highest under iron-replete conditions (R), though other model parameterizations capture canonical RyhB induction under iron limitation (Supplementary Figures S4 – S5). This pattern suggests that oxidative stress alone (RM) may mimic aspects of iron stress (D), while dual stress (DM) triggers a distinct or more robust regulatory response that overrides iron-conserving processes and reconfigures Fur-mediated control. Additionally, results suggest that RyhB may have a dual regulatory role, contributing to both iron homeostasis and the oxidative stress response (85).

The similarity between RM and D response extends beyond Fur and RyhB dynamics to iron transport variables (Supplementary Figures S5 – S6, SPMR ensemble results). Results show that ferrous uptake (feoB) is similarly induced under oxidative stress conditions (RM, DM), which have lower cell population relative to unstressed states (R, D). On a per-cell basis, ferrous uptake (feoB) shows stronger induction than ferric uptake (fepA) under oxidative stress, with both transporters maximally induced under dual stress (DM). Enterobactin dynamics in our model is consistent with previous findings (35); extracellular enterobactin (enter.e) peaked under deplete but is reduced in replete, while intracellular iron-bound enterobactin (feenter.c) is increased under oxidative stress (RM, DM). The synthesis of enterobactin in excess of transporter production suggests a secondary function for siderophores beyond iron chelation (35).

The model captures canonical Fe–S/IscR dynamics, with clusters increasing under replete conditions before plateauing, while IscR dips and recovers by autoregulation. Fe–S cluster pools (fes) are strongly depleted under D and DM, consistent with iron scarcity and vulnerability to oxidative stress. SPMR simulations (Supplementary Figures S5 – S6) instead show monotonic declines in Fe–S, a less biologically plausible response. ROS defenses in our model are significantly activated under exogenous, repetitive oxidant challenge with SodA (soda) having the greatest increase under dual stress (DM) conditions (48,49).

Biomass trajectories replicate experimental data and demonstrate how the complex interplay between stress response systems produces distinct physiological phenotypes. Cells in iron-sufficient media (R) grow exponentially, while iron-starved cells (D) exhibit bacteriostatic growth. Under oxidative stress, iron-replete cells (RM) appear to oscillate between bactericidal and bacteriostatic conditions, suggesting a bacteriostasis state where *E. coli* could resume growth with cessation of oxidative stress perturbations. However the added reduction in growth in dual stress (DM) conditions supports the bactericidal effect of combined iron and hydrogen peroxide stress (50). Collectively, our model results show that oxidant stress in iron-rich conditions (RM) yields metabolic trajectories resembling those of iron-starved cells (D) yet triggers a more adverse physiological response. These findings point toward a three-dimensional phenotype state space, with dynamic peroxide associated RM and DM forming a functional cluster distinct from replete (R) and deplete (D). Results also show how repeated stress (RM) facilitates phenotype plasticity, enabling *E. coli* to move between bacteriostatic and bactericidal outcomes (50).

### 2.8 Model Extensible to Quantify Differences in Siderophore Levels Under Novel Stress Condition

To evaluate the model’s robustness and extensibility beyond the original training and testing conditions, we used it to predict catechol-type siderophore production under dual-stress, replicating experiments conducted by Peralta et al. In their study *E. coli* BW25113 was grown for 20 hours in M9 medium under one of four conditions: (i) 1 mM H₂O₂, (ii) 25 µM FeCl₃, or (iii) neither or (iv) both. Siderophores not only scavenge iron but also buffer oxidative stress, making them an important, measurable functional outcome. Experimentally, exposure to peroxide alone increased catechol production by approximately 80% compared to control. We set our model’s initial iron pool and peroxide influx to reflect these perturbations and successfully reproduced the observed response hierarchy: maximal catechol output under H₂O₂ stress, intermediate levels under combined iron and peroxide, and minimal change with iron alone (Figure 8). Although absolute concentrations varied, due to differences in scaling and strain-specific parameters, the preservation of this ranking demonstrates the model’s capacity to generalize across stress conditions and bacterial strains, supporting its applicability beyond the original calibration dataset.

**Figure 8:**
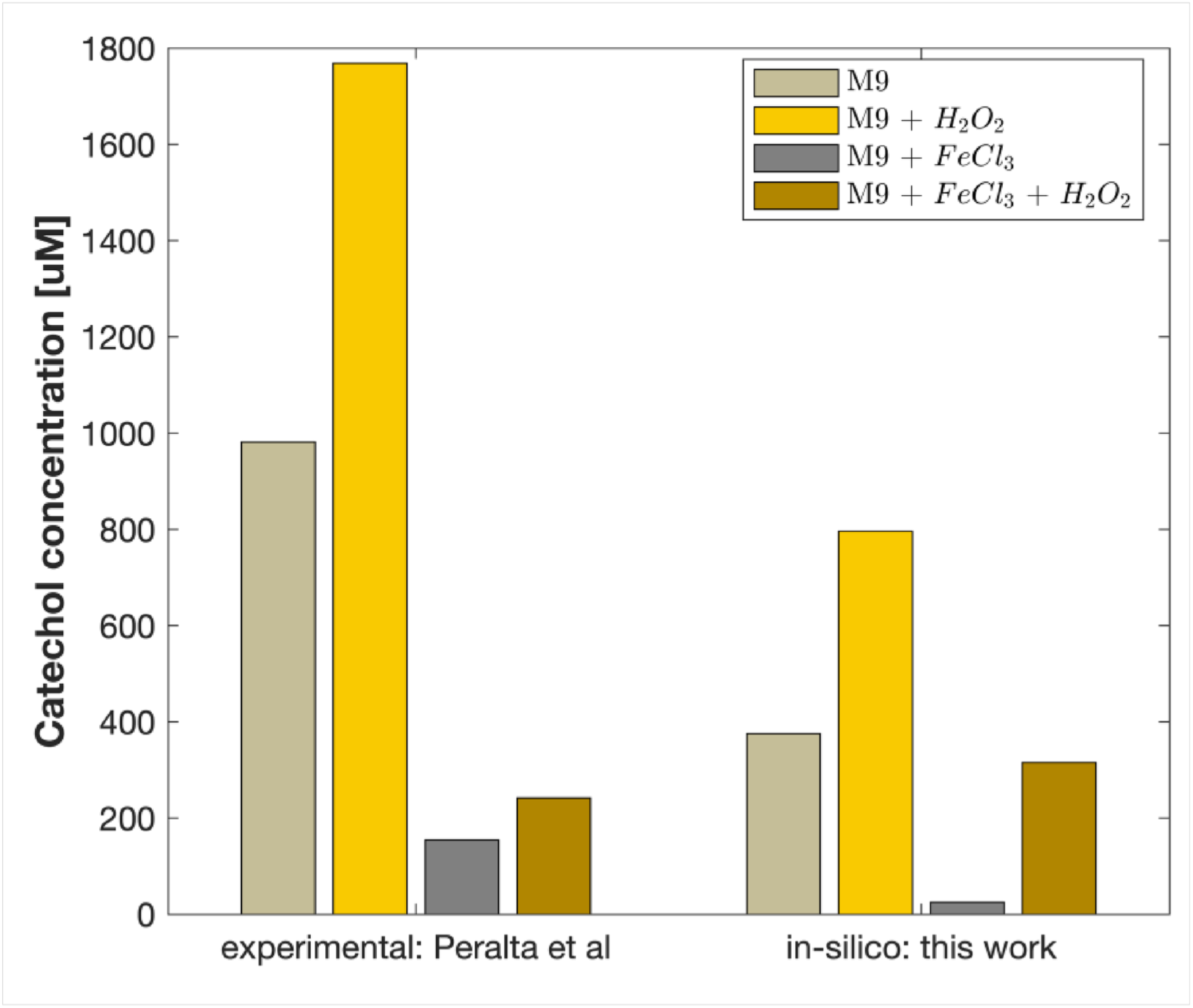
Catechol Concentration Comparison Under Various Stress Conditions.

### 2.9 Multi-Phenotype Optimization Enhances Physiological Relevance of Parameter Estimates and Reduces Suboptimal Parameter Entrenchment

We evaluated resulting model parameters to determine the relationship between in silico and empirically measured biochemical parameter ranges. The log-scaled marginal distributions for eight key parameters are compared across the three optimization schemes (Figure 9): maintenance cost (ksg), maximum growth rate (umax), Monod half constant (ks), catalase maximum velocity and Michaelis-Menten constant (Vmax and Km; vmaxcatr, kmh2o2kat), Fur–DNA affinity (kf), enterobactin esterase binding constant (kkenteres), and intracellular volume (vol.c). Empirical ranges from the BRENDA database are shown for comparison. Seven of the eight parameters remain within one order of magnitude of experiment for every optimization scheme, with MPMR models most comparable to empirical values for umax and ks.

**Figure 9:**
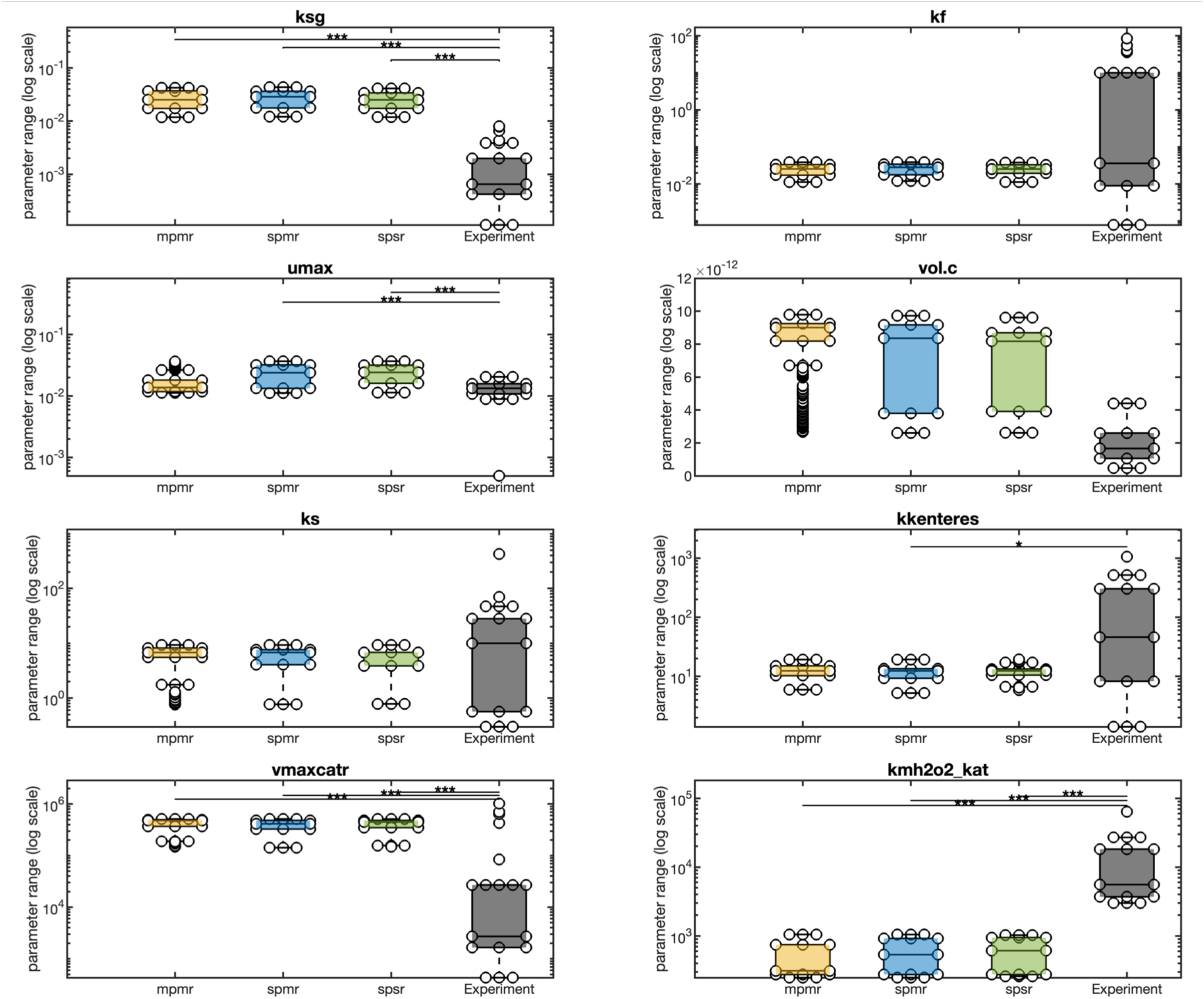
Parameter Space Analysis. From top to bottom (left to right), the figure compares log10 marginal distributions and the most probable values of growth maintenance factor and Fur:DNA dissociation constant (ksg and kf), iron induced growth constant and intracellular volume (umax and vol.c), Monod half constant due to iron and fes esterase hydrolysis rate (ks and kkenteres), Vmax and Km of catalase (vmaxcatr and kmh2o2kat) across the three schemes.

To determine how multi-phenotype and multi-response optimization affects parameter identifiability and stability, we analyzed the posterior distributions of all 35 calibrated model parameters, spanning Class I core kinetics and system-scale parameters, Class II regulatory parameters, and Class III subsystem and redox response parameters (Figure 10, Supplementary Figs. S7–S9). Using shape-based classification, each resulting distribution was assigned to one of six families reflecting differences in symmetry, tail behavior, and boundedness: Normal (N), Student’s t (t), Gamma (G), Lognormal (LN), Beta-U (Bu), and Beta-Bell (Bb).

**Figure 10:**
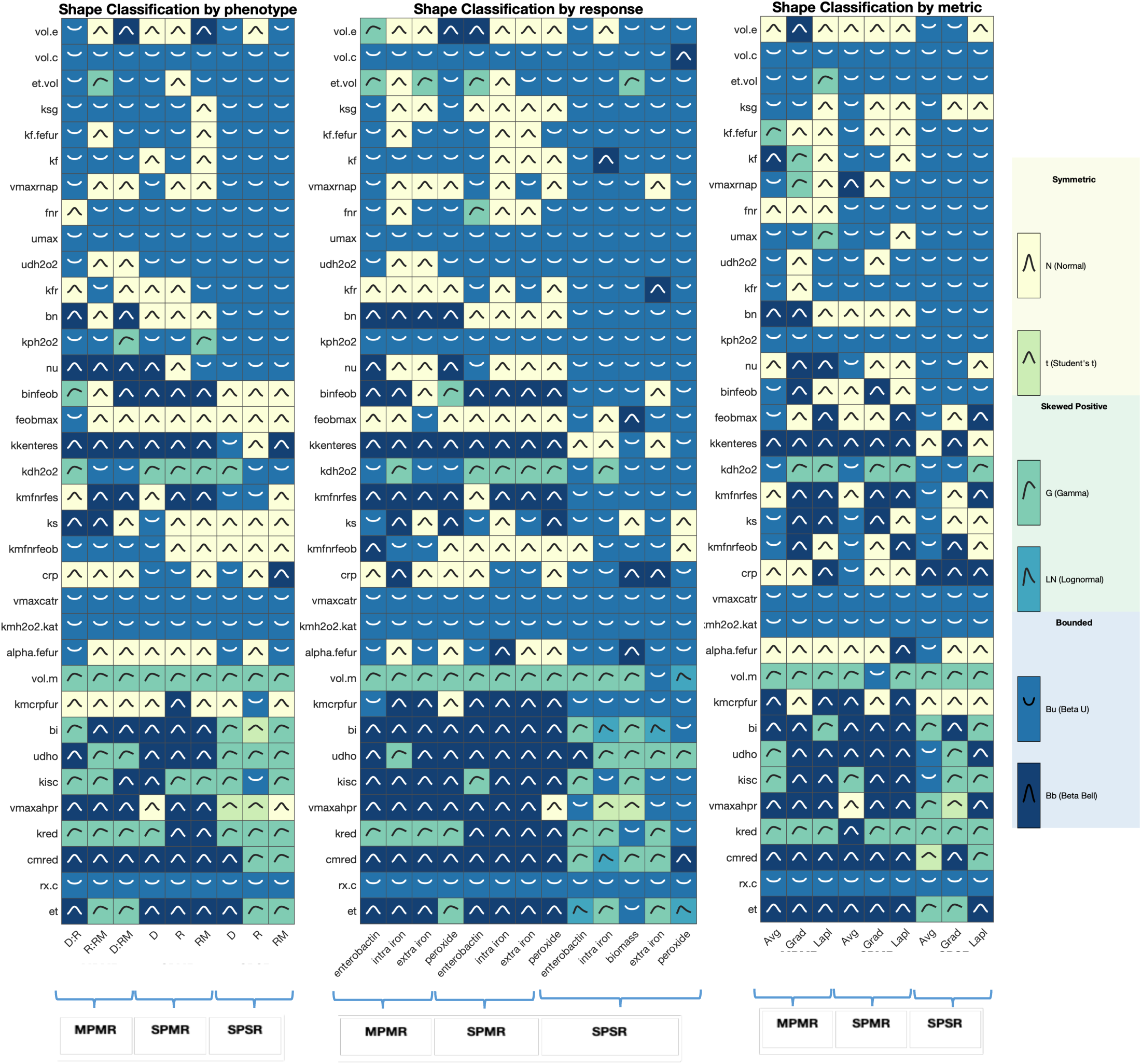
Classification of Posterior Parameter Distributions Across Optimization Strategies. The shape of posterior distributions was categorized for 35 optimized parameters (rows) across stress–response–metric combinations (columns), aggregated by (left to right) stress phenotype, response variable, and error metric. Each cell displays the determined shape class with an embedded, idealized distribution curve of the characteristic form. Parameter distributions were categorized into six shape classes based on symmetry, tail behavior, skewness, and boundedness: Normal (N), symmetric with light tails; Student’s t (t), symmetric with heavy tails; Gamma (G), right-skewed with moderate tails; Log-normal (LN), right-skewed with heavy tails; Beta-U (Bu), bounded with U-shaped density; and Beta-Bell (Bb), bounded with bell-shaped density. Related distribution families are grouped by color: symmetric distributions (N, t) yellow tones; positively skewed distributions (G, LN), green tones; bounded distributions (Bu, Bb), blue tones. Across optimization strategies, distribution shapes were inferred from 4,968 parameter sets in MPMR (36 combinations × 138 models), 4,896 parameter sets in SPMR (36 × 136), and 135 parameter sets in SPSR (45 × 3). Where subsampling was required for comparability, analyses were performed using the top 135 models, selected by composite score, coverage, or error, as appropriate. **Parameter abbreviations:** vol_e, extracellular volume; vol_c, intracellular volume; et_vol, volume transfer coefficient between extracellular and intracellular compartments; ksg, growth maintenance factor; kf_fefur, Fur–Fe²⁺ association constant; kf, Fur–DNA dissociation constant; vmaxrnap, maximal RNA polymerase transcription rate; fnr, Fnr transcription factor concentration or activity; umax, iron-induced growth constant; udh2o2, peroxide-induced death constant; kfr, nutrient conversion factor; bn, iron-starvation scaling factor; kph2o2, metabolic hydrogen peroxide production rate; nu, Fe²⁺ to Fe³⁺ nutrient conversion ratio; binfeob, FeoB-mediated iron uptake rate in a fur mutant; feobmax, maximal cellular FeoB transcription rate; kkenteres, Fes esterase hydrolysis rate constant; kdh2o2, Monod peroxide growth half-saturation constant; kmfnrfes, Fnr–DNA dissociation constant for the fes promoter; ks, Monod half-saturation constant for iron; kmfnrfeob, half-saturation constant for Fnr-dependent FeoB transcription; crp, CRP (cAMP receptor protein) concentration; vmaxcatr, maximal catalase reaction rate; kmh2o2_kat, catalase half-saturation constant for H₂O₂; alpha_fefur, Fur–Fe²⁺ binding cooperativity factor; vol_m, extracellular volume occupied per single cell in the medium; kmcrpfur, CRP–DNA dissociation constant for the fur promoter; bi, scaling factor for Isc Fe–S cluster synthesis; udho, maximal growth-rate reduction coefficient due to hydroxyl radicals; kisc, saturation constant for Isc Fe–S cluster assembly; vmaxahpr, maximal alkyl-hydroperoxidase reaction velocity; kred, reductase production rate; cmred, half-maximal inhibitory peroxide concentration for reductase activity; rx_c, intracellular concentration of reducing equivalents; and et, enzyme scaling factor.

Across optimization strategies or modes, systematic shifts in distribution families were observed as dataset dimensionality increased. The prevalence of Beta-U distributions decreases monotonically from SPSR (63.6) to multi-response SPMR and multi-phenotype MPMR ( 34.3% - 35.1%), while Beta-Bell distributions increase from ∼7% in SPSR to ∼27% in SPMR and MPMR. Normal distributions are most prevalent in SPMR (∼28%) and MPMR (∼23%). These differences indicate a systematic reduction in boundary-based parameter assignments and a more robust exploration of the parameter space under multi-response and multi-phenotype methods.

Parameter classes, optimized over three sequential steps, differed in distribution shape. Sequence I parameters (Class I) are dominated by Beta-U distributions (66.6%) with a Bu:Bb ratio of 11:1, indicating persistent boundary concentration regardless of optimization mode. Sequence II parameters (Class II) show intermediate behavior with Bu:Bb ratio of 2:1. Most notably, Sequence III parameters (Class III) have the inverse composition, with Beta-Bell distributions dominating (43.1%) over Beta-U (18.1%), yielding a Bu:Bb ratio of 1:2. This progression suggests that these locally acting parameters benefit most from multi-response optimization, resulting in centrally-concentrated posteriors.

Distribution homogeneity, the degree to which a parameter retains the same distribution family across evaluated strategies, is indicative of how optimization strategies differentially constrain parameter space. Each of the three optimization methods produce a small core group of approximately nine to ten parameters that maintain the same distribution family across stress phenotype, response measures, metric, or all three subgroupings. Under SPSR, all nine homogeneous parameters (kf_fefur, fnr, umax, udh2o2, bn, kph2o2, vmaxcatr, kmh2o2_kat, and rx_c) are classified as Beta-U, indicating accumulation of optimal parameter values near the boundaries regardless of dataset organization. In contrast, homogeneous parameters in MPMR span multiple distribution families, including Beta-U (vol_c, vmaxcatr, kmh2o2_kat, rx_c) and Beta-Bell (kkenteres, vmaxahpr, cmred), and Gamma (vol_m, kred). SPMR exhibits a similar diversity with homogeneous parameters distributed across Beta-U, Beta-Bell, and Normal families.

Distribution heterogeneity, the degree to which parameters switch across distribution families provides insight into context-sensitivity to phenotypes and measures, or instability. SPSR displays the greatest heterogeneity in its Class III (mostly redox) parameters, with posteriors for bi and vmaxahpr spanning up to five distinct distribution families. Symmetric heavy-tailed (Student’s t) and right-skewed heavy-tailed (Lognormal) distributions appear exclusively in SPSR suggesting single-response optimization includes tail-equivalent solutions where substantially different parameter values selected under different phenotype or response conditions achieve comparable fit. Under MPMR, this heterogeneity is largely absent for the same subsystem parameters, with vmaxahpr, kred, and cmred classified predominantly as Beta-Bell or Gamma distributions; this demonstrates how additional phenotypes and responses effectively constrains the solution space to a subset of comparably distributed values, thereby reducing ambiguities and instability of multi-family switching observed in SPSR.

Overall parameter distribution patterns show that SPSR is characterized by widespread boundary concentration, occasional heavy tails, and tends towards extreme boundary values, which is consistent with practical non-identifiability challenges. Whereas SPMR and MPMR are associated with reduced or narrowed boundaries, elimination of heavy-tailed families, and increased stability of posterior distributions across optimization conditions. Results support that multi-phenotype and multi-response optimization provides stronger parameter constraints, yielding a more stable posterior structure enabling model robustness and extensibility.

## 3.0 Conclusion

In this work we constructed a mechanistic model to predict bacterial response to iron and oxidative stress and developed a novel, extensible multi-phenotype, multi-response optimization framework for parameterization of robust models for dynamic, multi-stress environments. We used our framework to investigate *E. coli* behavior across four iron-peroxide response phenotypes, combining experimental time-series datasets, augmented sensitivity analysis, and sequential (“identify-then-refine”) optimization workflows to fit broadly influential to more limited impact parameters. Model robustness of candidate models was improved using a weighting-based selection algorithm and linear programming to construct an optimal ensemble model that minimized overall prediction error. This strategy produced a 50-variable ODE model that (i) recapitulated the dynamics of growth, iron pools, peroxide dismutation, and siderophore secretion under each stress combination, (ii) demonstrated enterobactin contribution to both iron scavenging and oxidative-stress mitigation, and (iii) predicted qualitative shifts in Fur activation, Suf/Isc transisiton, and *RyhB* suppression that mirror empirical observations.

Biologically, the simulations show that repeated peroxide stress reduces the Isc Fe-S assembly pathway while maintaining the peroxide-resistant Suf system, with holo-Fur moderately increased (Figure 11). In dual-stress conditions, intracellular Fe-enterobactin accumulates substantially while extracellular Fe-enterobactin increases modestly, suggesting repeated peroxide promotes iron chelation and intracellular retention rather than blocking export. Peroxide stress also produces elevated SodA levels (increased under iron-depleted dual stress), near-bacteriostatic growth under dual and single stress conditions (average biomass fold-change reduced under iron deplete and iron-replete conditions), and it correctly ranks catechol siderophore output highest with iron depletion and lowest when iron is replete, confirming enterobactin’s role as a redox buffer.

**Figure 11.**
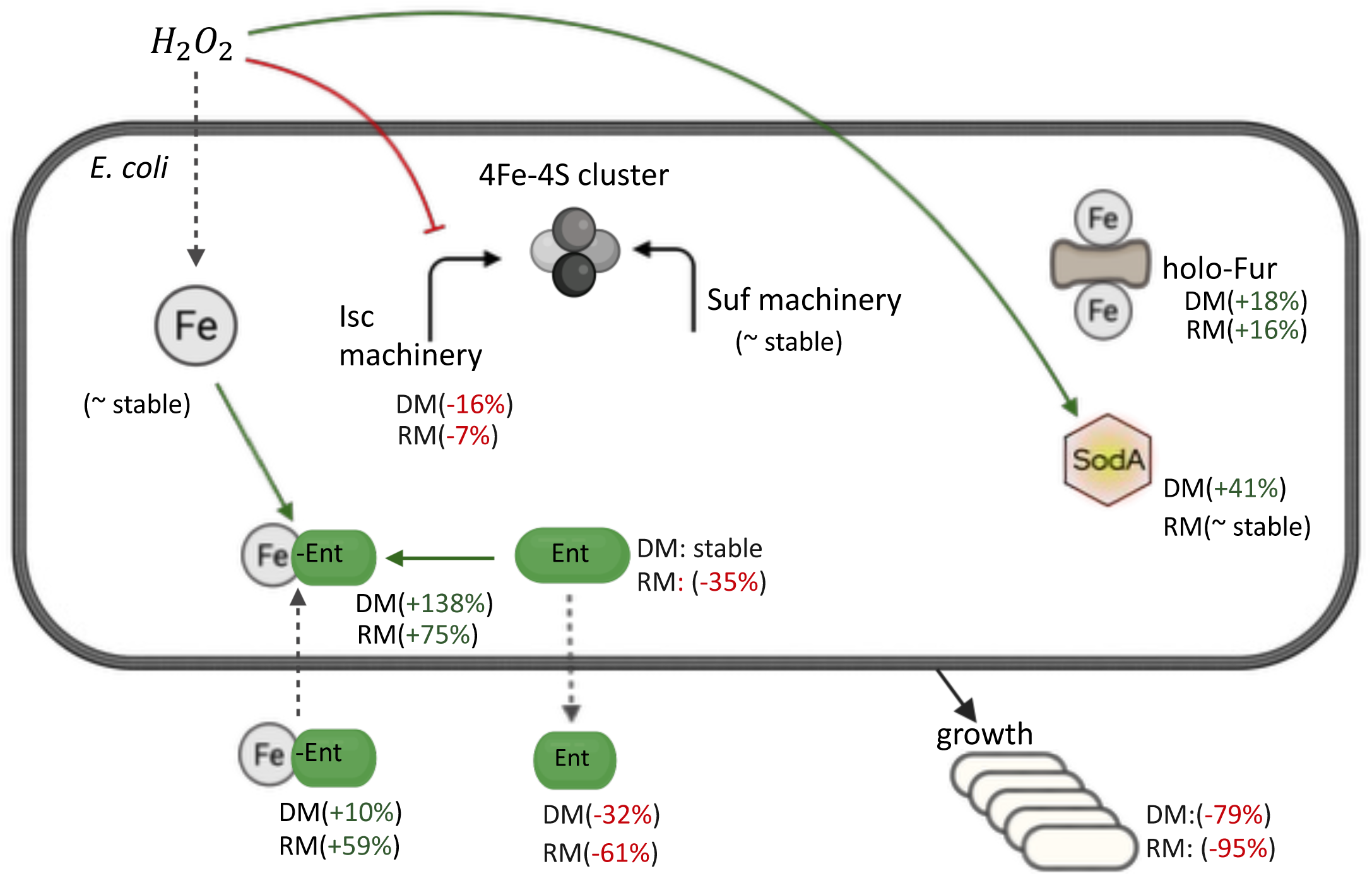
Summary of model-derived insight on E. coli response to single and dual (H₂O₂ + iron limitation) dynamic peroxide stress. The effects of repeated peroxide stress under iron-deficient and sufficient conditions on E. coli iron homeostasis, oxidative stress response, and growth are represented (summary of model outcomes from Supplementary Tables S3 and S4). Components include cytoplasmic Fe²⁺, the Isc and Suf Fe-S assembly pathways, 4Fe-4S clusters, iron-sensing regulator holo-Fur, antioxidant enzyme SodA, enterobactin (Ent) and Fe-enterobactin (Fe-Ent) in both intracellular and extracellular compartments, and growth (biomass). Line types: grey dashed, transport; red, inhibition; green, activation. Percentages indicate changes under dual stress (DM) or peroxide stress (RM) where indicated.

Beyond demonstrating mechanistic fidelity and biological insight, we also assessed how well our fitted parameters align with independent empirical benchmarks. Importantly, our multi-phenotype and multi-response calibration not only increased coverage (∼80% vs 60%) but also produced biologically realistic parameter estimates, with multi-phenotype calibration estimates within one order of magnitude of published empirical ranges for five of eight key parameters. Analysis of posterior distributions demonstrated that optimization approach fundamentally shapes parameter identifiability. Single-response optimization (SPSR) yielded distributions dominated by boundary-concentrated Beta-U shapes, indicative of parameter values at the extreme and consistent with practical non-identifiability. In contrast, multi-response optimization strategies (SPMR and MPMR) produced a balanced mix of bounded distributions (Beta-U and Beta bells), with heavy-tailed distribution families (Student’s t, Lognormal) nearly eliminated. Strikingly, parametric class was as an important if not more important determinant of distribution form. Distributions of core kinetic parameters (Class/Sequence I) remained boundary-concentrated regardless of optimization approach, while redox subsystem parameters (Class/Sequence III) under multi-response optimization, shifted from boundary- to centrally-concentrated posterior distributions. This suggests that these locally acting parameters benefit disproportionately from multi-phenotype, multi-response calibration methods.

Results of our study produced three key insights. First, capturing system dynamics across divergent phenotypes requires data diversity more than data volume; the MPMR pipelines consistently yielded the most robust parameter sets and the most stable posterior distributions. Second, post optimization approaches such as error weighting offers a practical hedge against phenotype-specific bias as measured by using error biasing to improve single phenotype single response models while maintaining their mechanistic interpretability. Third, even well-behaved global fits can underperform at capturing transient iron–ROS couplings under dual-stress scenarios as revealed in our multi-phenotype, multi-response analysis, which suggests structural gaps that targeted model refinement could address. Additionally, future improvements can strengthen our current multi-phenotype, multi-response optimization framework. First, integrating formal uncertainty quantification into the calibration loop would provide confidence bounds on both parameter estimates and model predictions particularly for Class III redox parameters where posterior stability improved most dramatically under MPMR. Second, hybrid optimization schemes could improve convergence speed by using machine-learning surrogates to rapidly identify candidate parameters that can be refined using mechanistic models to simulate and evaluate biological realism. Third, targeted refinements to the model structure, particularly to subsystems involved in iron–ROS coupling under dual stress, would help close the gaps we observed in transient behaviors. Finally, extending the calibration to additional *E. coli* strains or related species under varied environmental contexts would test the model’s robustness, extensibility, and applicability beyond MG1655 to guide both experimental design and therapeutic strategies in iron- and peroxide-stressed settings.

## 4. Methods

### 4.1 Experimental Multi-stress Response Model

The empirical data used in this study was generated from a time-resolved multi-stress experiment carried out with *Escherichia coli* MG1655 (ATCC 700926) grown in M9 minimal media (19, 53). Briefly, iron availability was manipulated by treating the base media (M9 Complete) overnight Chelex^®^ 100 Chelating Resin (BioRad). The resulting iron-depleted media (∼ 0.1 µM residual Fe) served as the deplete (D) condition, while supplementation with 7.2 µM FeSO₄ defined the replete (R) condition. One liter cultures were inoculated from 24 h overnight cultures (OD₆₀₀ ∼0.04) and grown at 37°C for 12 hours. Starting post-inoculation (0 hr) and every 90 min thereafter, Milli-Q water or 100 µM H₂O₂ was added (spiked) to cultures resulting in four environments replicated in our computational model: iron-replete (R), iron-deplete (D), iron-replete plus peroxide (RM), and iron-deplete plus peroxide (DM). Cultures were sampled before and 30 minutes following water or H₂O₂ spiking and used to quantify several responses, including: biomass via colony forming units (log CFU mL⁻¹ by plating), intracellular iron (*KMnO₄*-ferrozine protocol), and colorimetric assays to quantify extracellular ferrous iron, hydrogen peroxide (Abcam), and catecholate-type siderophores (Emergen Bio). Data interpolation using the aggregate trend across two biological replicates was used to fill in missing values. Our model uses the post-spike data to represent *E. coli’s* response to perturbation, which was categorized into metabolic, environmental, and growth response measures under iron and oxidative stress condition and used to calibrate the integrated iron-peroxide model described.

### 4.2 Theoretical and Computational Model Development

#### 4.2.1 Modeling the *E. coli* Iron Regulatory Network

At the core of the *E. coli* iron regulatory network is the ferric uptake regulator protein (Fur), whose activity depends on its binding to Fe²⁺. This iron cofactor requirement is central to Fur’s regulatory logic as depicted in Figure 12. Under iron-sufficient conditions, the active Fur-Fe²⁺ complex represses genes involved in iron uptake and suppresses the expression of the small RNA RyhB, which normally downregulates non-essential iron protein production. This dual repression prevents excess iron accumulation and ensures that iron is allocated to essential cellular functions. When iron levels decline, Fur becomes inactive, permitting increased expression of RyhB. Elevated RyhB then targets mRNAs encoding non-essential iron proteins for degradation, thereby redirecting available iron to support essential functions (1,11,54). In addition to managing iron uptake and utilization, cellular iron function depends on the assembly of iron-sulfur (Fe-S) clusters. In *E. coli*, this process is controlled by two main systems: the housekeeping Isc pathway and the stress-activated Suf pathway (52). The transcriptional regulator IscR plays a critical role in maintaining the Fe–S pool by employing an autoregulatory feedback mechanism through its binding to Fe–S clusters (55). Under iron-sufficient conditions, Fur maintains expression of the Isc machinery by repressing RyhB and directly repressing the stress-induced suf operon. When iron is scarce or peroxide stress oxidizes Fe²⁺ to Fe³⁺, destabilizing Fe–S cluster assembly, Fur is inactivated, which derepresses both RyhB and the Suf system. In response, IscR senses the diminished Fe–S pool and modulates isc operon expression accordingly. This integrated regulation allows *E. coli* to balance iron acquisition, Fe-S cluster assembly, and iron utilization, thereby protecting the cell from both iron deficiency and oxidative damage.

**Figure 12:**
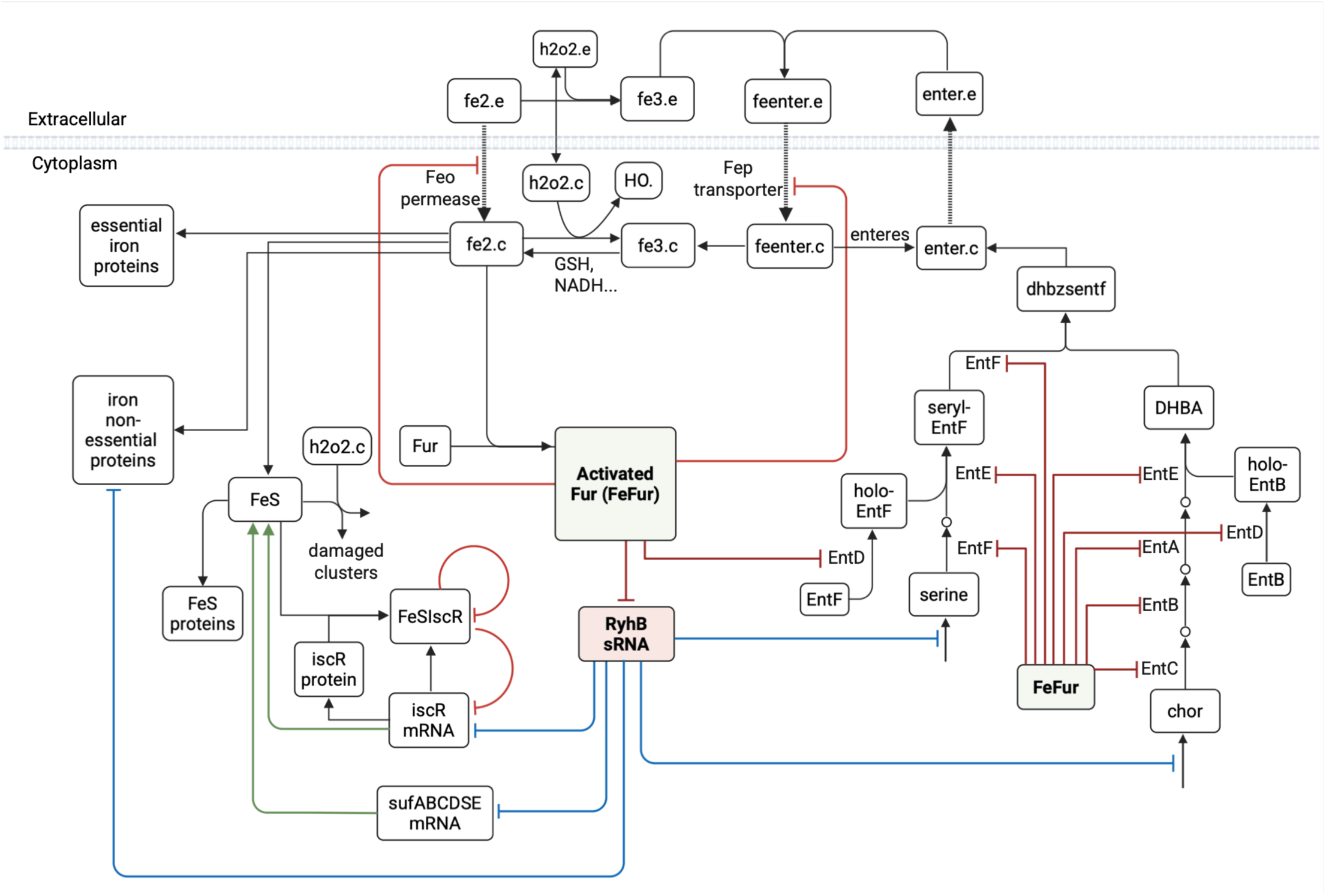
Summary Diagram of Iron and Peroxide Stress Response Network in E. coli. Schematic of the integrated iron regulatory network model showing iron uptake (Feo permease for Fe²⁺; Fep transporter for ferric-enterobactin), the central Fur-Fe²⁺ regulatory hub, RyhB-mediated post-transcriptional control, Fe-S cluster homeostasis via IscR and the Suf system, and the enterobactin biosynthesis pathway. Green lines indicate transcriptional activation; Red lines indicate repression; blue lines indicate post-transcriptional control; black arrows denote biochemical reactions or transport. Species that occur across compartments are delineated ‘.c’ and ‘.e’ representing cytoplasmic and extracellular species respectively.

#### 4.2.2 Modeling Iron Acquisition in *E. coli*

*E. coli* uses a range of transporters that fall in two broad iron acquisition categories: diffusion versus siderophore mediated. Ferrous iron (Fe²⁺) uptake occurs via diffusion through specific permeases. In our model, this process is primarily represented by the Feo system, encoded by the feoABC operon and regulated by Fur-Fe²⁺ and Fnr (3, 56). In contrast, ferric iron (Fe³⁺) is acquired through siderophore-mediated mechanisms. Although *E. coli* releases several siderophores under iron-deficient conditions, our model focuses on enterobactin, the prototypical catecholate siderophore with the highest known affinity for Fe³⁺ (57). Enterobactin binds extracellular Fe³⁺ to form ferric-enterobactin complexes, which are then imported into the cell via specific transport proteins, such as those encoded by the fep operon, a major transporter for catecholate-type siderophores.

##### 4.2.2.1 Enterobactin Biosynthesis and Export

Enterobactin biosynthesis in *E. coli* occurs in two stages: first, chorismate (CHOR) is converted to 2,3- dihydrobenzoate (DHB) and then enterobactin (ENTER) is synthesized through the stepwise ligation of DHB and L-serine (SER) (54,58). While the pathway has been extensively characterized (58–65), its dynamic regulation in response to iron and oxidative stress and in particular its contribution to iron regulation has not been modeled mechanistically. To close this gap, we developed a multi-tiered family of models representing the biosynthetic pathway at increasing levels of resolution (Supplementary Materials Appendix S5, Table S1) to determine the model that best captures the iron regulation and control. Figure 13 shows the enterobactin biosynthesis model used, highlighting the multi-step transcriptional regulation by Fur that drives the bacterial dynamic response to iron limitation and oxidative stress.

**Figure 13:**
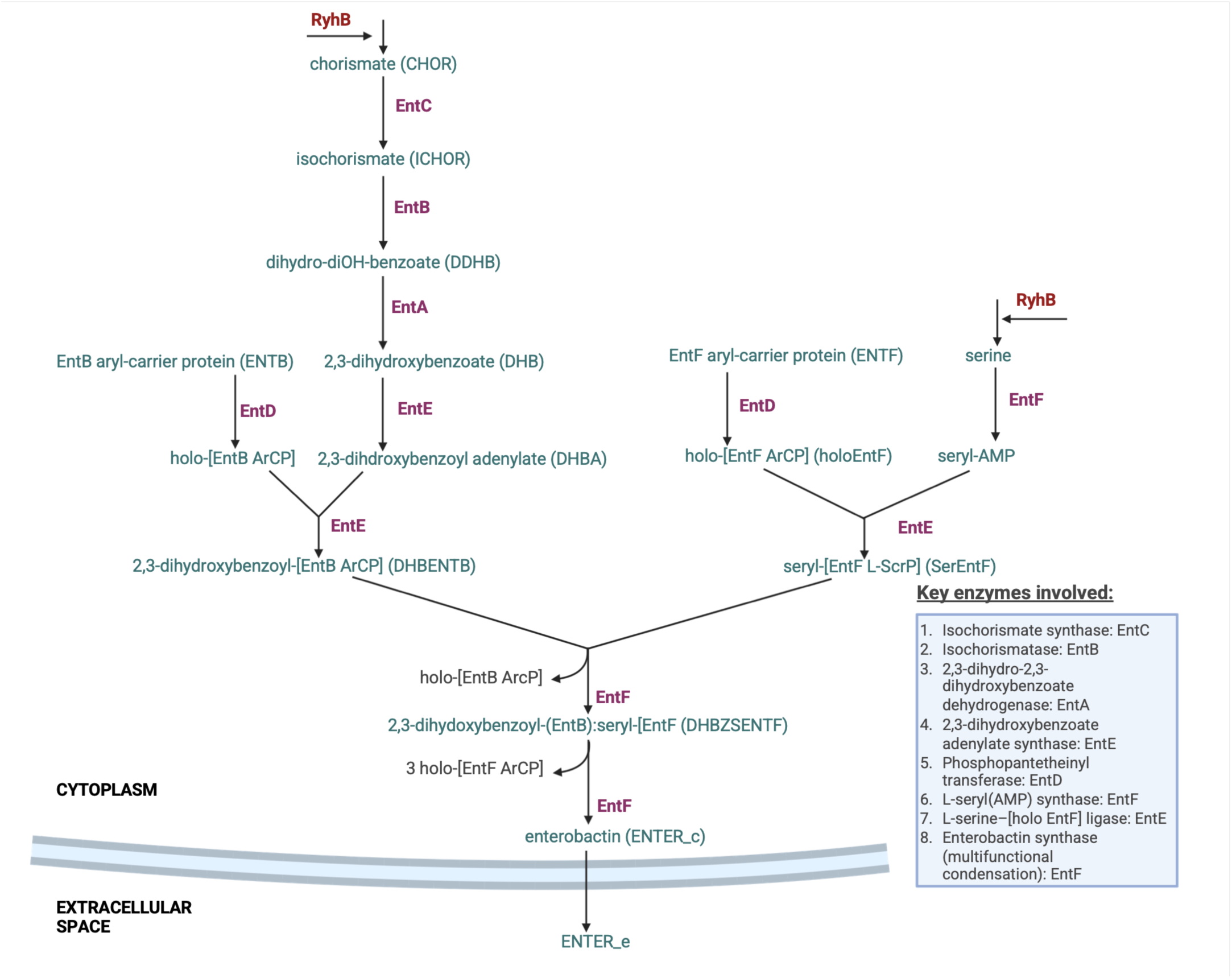
Enterobactin Biosynthesis Network in E. coli. Schematic of the two-stage enterobactin biosynthesis model. The left branch depicts conversion of chorismate to 2,3-dihydroxybenzoyl adenylate (DHBA) via EntC, EntB, EntA, and EntE, with holo-EntB formation by EntD. The right branch shows L-serine adenylation and loading onto holo-EntF. Committed reactions at both branches are regulated by the small RNA RyhB under iron limitation. Convergence at the DHBZSENTF intermediate leads to EntF-catalyzed assembly of cyclic enterobactin (ENTER_c), which is exported extracellularly (ENTER_e). Key enzymes are listed in the inset box.

Enterobactin biosynthesis begins with chorismate (CHOR), which is a product of the shikimate pathway (58). Chorismate dynamics is described as

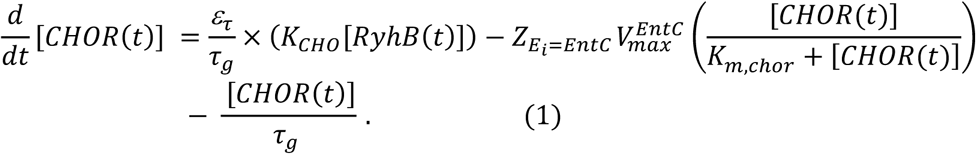

The first term in Equation 1 represents production, which depends on *RyhB* concentration and ε_*τ*_, the enzyme scaling factor for the siderophore associated biosynthesis pathways. *RyhB* channels metabolism towards the enterobactin biosynthesis pathway via the shikimate permease *shiA* under iron stress conditions (9, 57, 59). To capture this regulatory effect, we use a basal production rate for chorismate *k_CHO_* that is modulated by the cellular level of *RyhB*, thereby actively incorporating RyhB-mediated permease activation without explicitly modeling the downstream components of the shikimate pathway. The second term represents the consumption of chorismate through its conversion to isochorismate by EntC, modulated by the Fur regulatory factor Z (see Equation 2).

As in the model by Semsey et al., basal production and usage (third term) are scaled by a dilution rate *τ*_*g*_ to maintain a consistent intracellular concentration over time and account for dilution of chorismate due to cell division.

In contrast to reactions indirectly regulated by Fur via *RyhB*, enzymes involved in intermediate enterobactin biosynthetic reactions are directly regulated by Fur and are modeled using Michaelis-Menten kinetics. *Fur-Fe^2^* regulation is incorporated by modulating each enzyme’s *V_max_* value with a gene-associated regulatory factor *Z*_*E_i_*_, (Equation 2) (66 - 67,105). Consistent with this approach, we explicitly model the bifunctional proteins EntB and EntF, which serve as both upstream substrates and downstream enzymes, while EntC, EntD, EntE, and EntA are not explicitly modeled. However, their transcriptional regulation by Fur is captured via the regulatory term *Z*_*E_i_*_. The first stage in the biosynthetic pathway captures the sequential conversion of chorismate (CHO) to isochorismate (ICHO), ICHO to 2,3-dihydroxy-2,3-dihydrobenzoate (DDHB), DDHB to 2,3-dihydroxybenzoate (DHB), and finally DHB to 2,3-dihydroxybenzoyl adenylate (DHB-AMP). The rate of change of the *i^th^* metabolite in these steps is modeled as

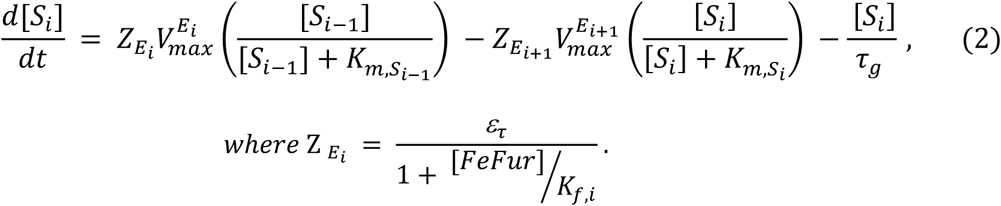

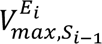 is the maximal velocity of the *i^th^* enzyme, *E_i_* acting on precursor metabolite *S*_*i*–1_. *K*_*m*,*S*_*i*−1__ is the Michaelis-Menten constant for the *i^th^* enzyme specific to substrate *S*_*i*–1_. *Z*_*E_i_*_ is a gene-associated modulation factor that conveys Fur’s transcriptional regulation on enzyme *E*_*i*_, representing regulatory effects based on transcription factor binding kinetics at the enzyme’s promoter (66, 105). In the presence of iron, *Fur-Fe^2+^* inhibits the production of the *(i-1) ^th^* enzyme acting on the *i^th^* substrate while the converse is true in the absence of iron. *K*_*f*,*i*_ denotes the dissociation constant for *Fur-Fe^2+^* complex at the regulatory site of the enzyme that produces the *i^th^* pathway intermediate. In our simulations, we initialized each *K*_*f*,*i*_ to 7 × 10^-6^*uM* (68).

In the second stage of enterobactin synthesis, 2,3-DHB is combined with L-serine to form enterobactin. In our model, this process occurs over seven reaction steps, beginning with serine adenylation by enterobactin synthase (*EntF*). Equation 3 describes serine dynamics in our model.

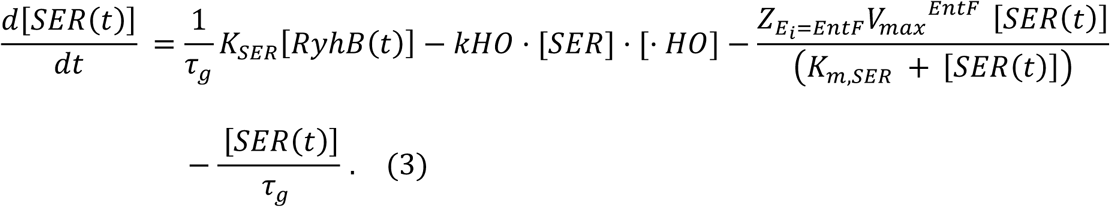

The first term captures serine synthesis upregulation by RyhB. By repressing cysE translation, thus lowering serine acetyltransferase activity for cysteine production, RyhB redirects excess serine into the enterobactin pathway (56,69). This is captured by multiplying the basal serine production rate *k_SER_* by the RyhB level and scaling by a dilution rate *τ*_*g*_ to maintain a consistent intracellular concentration over time. The second term represents hydroxyl radical damage to serine (see Equation 26). The third term represents serine adenylation by the L-seryl ligase, EntF, to produce (L-seryl)-adenylate (SERAMP). The fourth term accounts for growth associated dilution of serine.

The dynamics of (L-seryl) adenylate (SERAMP) is described as (Equation 4):

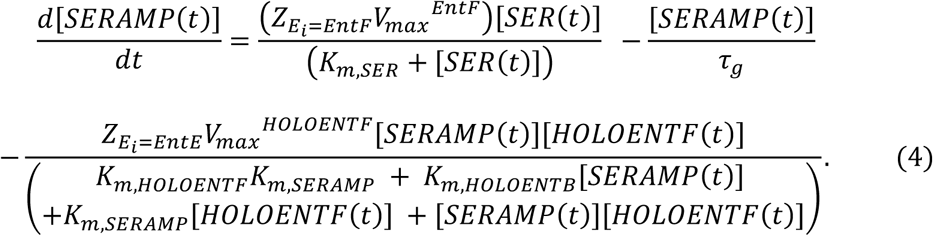

The first term represents production via EntF-catalyzed adenylation of serine, modulated by the factor *Z*_*E*_*i*_=*EntF*_. The second term accounts for growth associated dilution of seryl-adenylate during cell division. The third term represents SERAMP dependent synthesis of SerEntF via EntE’s ligase activity (holo-EntF ligase).

The dynamics of holo-EntB formation can be expressed as (Equation 5):

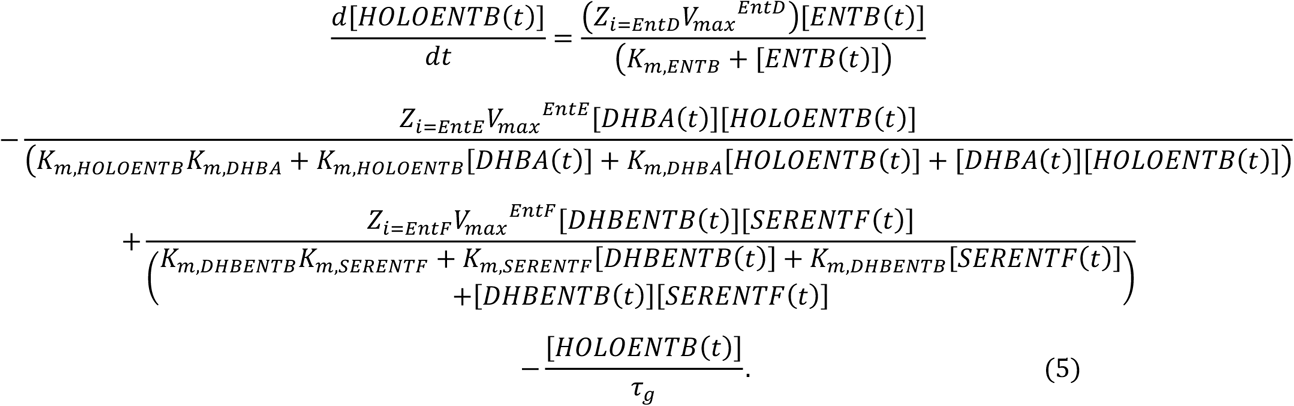

The first term represents the conversion of apo-EntB to holo-EntB by phosphopantetheinyl transferase EntD, modulated by the regulatory factor, *Z*_*i*=*EntD*_. The second term captures the EntE-mediated ligation of holo-EntB with DHB-AMP to produce 2,3-DHB-EntB. The third term accounts for a subsequent reaction step (described in Equation 9) that produces holo-EntB, and the fourth term represents dilution due to cell growth.

EntD also catalyzes the phosphopantetheinylation of apo-EntF, and the dynamics of this reaction are expressed as follows (Equation 6):

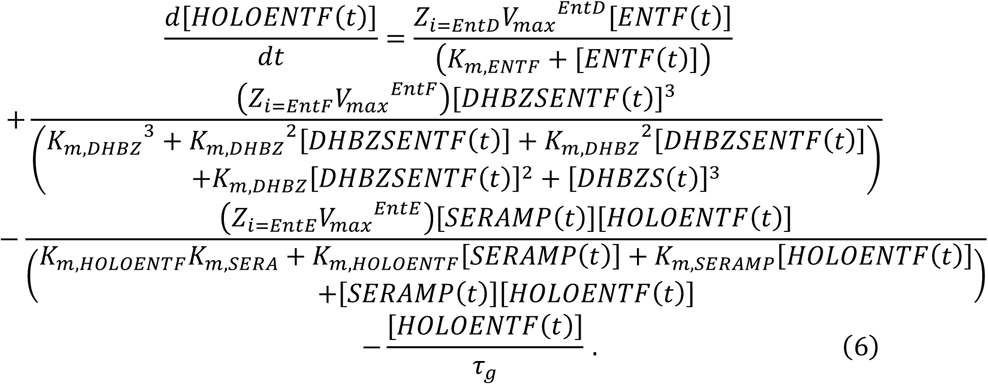

The first term represents the conversion of apo-EntF to holo-EntF by phosphopantetheinyl transferase EntD. The second term captures the regeneration of holo-EntF during enterobactin cyclization, where three molecules of DHBZSENTF are consumed to form cyclic enterobactin (see Equation 10). The third term describes the ligation of SerAMP with holo-EntF by the L-serine–[holoEntF] ligase EntE to yield SerEntF. The fourth term accounts for dilution of holo-EntF due to cell division.

The dynamics of *SerEntF* formed from the ligation reaction is described as (Equation 7):

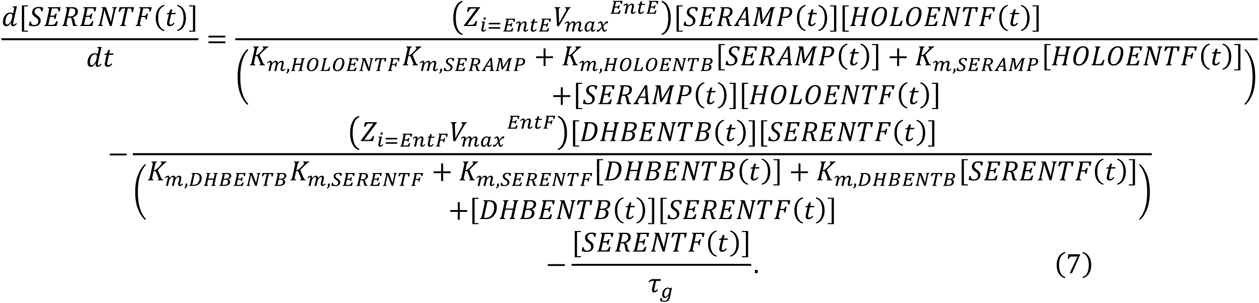

The first term represents the production of SerEntF from the ligation reaction between SerAMP and holo-EntF, catalyzed by EntE and modulated by its regulatory factor *Z*_*i*=*EntE*_. The second term captures the consumption of SerEntF during its ligation with DHB–EntB to yield the DHBZS intermediate, a necessary step toward enterobactin synthesis. Finally, the third term is a dilution term that accounts for the reduction in SerEntF concentration due to cell division.

SerEntF formed from the ligation reaction between SerAMP and holo-EntF undergoes two successive linkage reactions with DHB to yield enterobactin. Upstream, EntE first transfers the DHB acyl group onto holo-EntB, yielding acylated EntB (DHB-EntB, Equation 8). This is followed by the ligation of SerEntF with DHB-EntB, as represented in Equation 7, which is necessary for the formation of 2,3-dihydroxybenzoyl-[2,3-dihydroxybenzoyl-carrier protein] (DHBZS, Equation 9).

Equation 8 describes the kinetics of DHB-EntB formation.

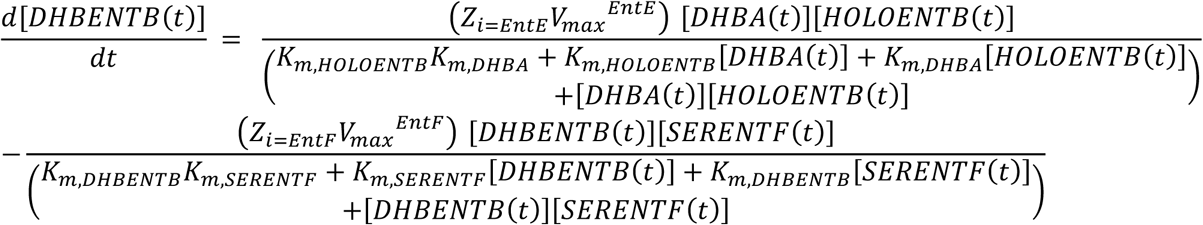

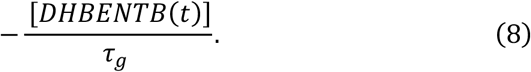

The first term represents the ligation of activated DHB with holo-EntB, mediated by EntE and modulated by the enzyme regulatory factor. The second term accounts for the consumption of DHB-EntB in the subsequent linkage reaction with SerEntF. The final term represents dilution of DHB-EntB due to cell growth.

The dynamics of *DHBZSENTF* formation is described as (Equation 9):

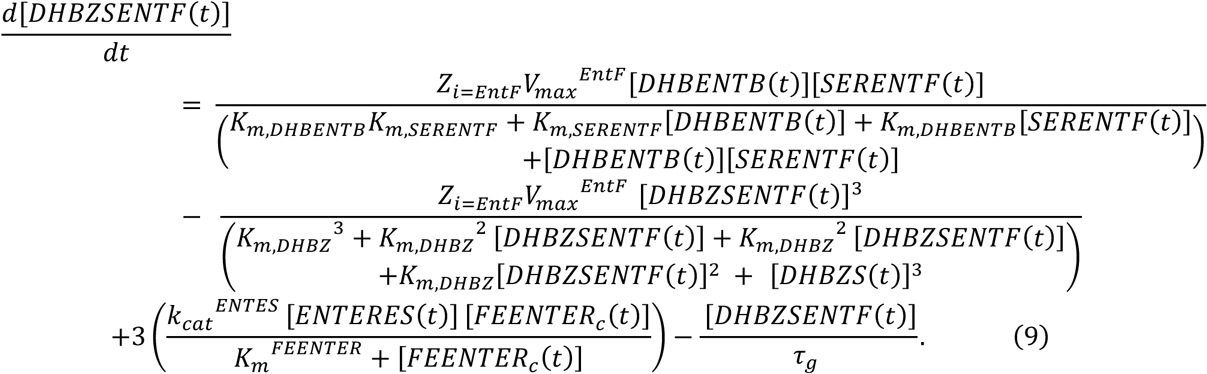

In our model, EntF catalyzes the assembly of enterobactin via multiple linkage reactions. The first term represents the EntF-mediated ligation of DHB-EntB with SerEntF, during which a molecule of holo-EntB is released. The second term captures EntF’s role in catalyzing the formation of the amide and ester linkages necessary for cyclic enterobactin synthesis. The third term reflects the hydrolysis of the ferric-enterobactin complex, releasing three molecules of DHBZS, and the final term accounts for dilution due to cell division. This formulation captures the critical enzymatic steps underlying enterobactin biosynthesis in response to dynamic iron stress.

In the second stage, enterobactin is synthesized by EntF through the formation of three ester linkages between three N-acylated serine residues. Equation 10 describes this process:

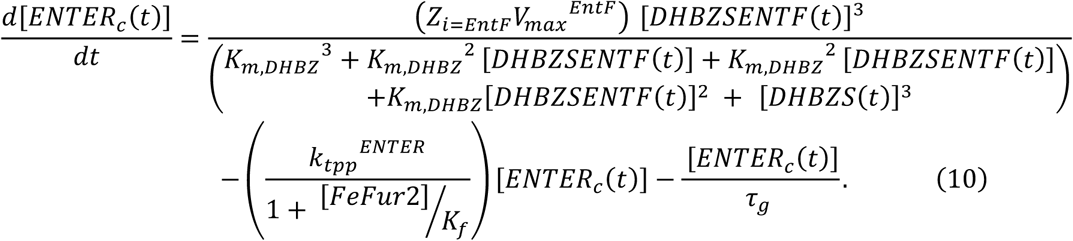

The first term represents the cooperative conversion of DHBZSENTF into cyclic enterobactin via EntF-catalyzed reactions following Michaelis–Menten kinetics. The second term models the export of enterobactin using a first-order kinetic model, an export process that although not fully elucidated is thought to be mediated by EntS via a Tol-C efflux system and is regulated by Fur-Fe²⁺(69 - 71). The final term accounts for dilution due to cell division.

#### 4.2.3 Modeling Gene Expression and Protein Translation in the Iron Regulatory Network

We represent protein production as a two-step process, transcription and translation, and capture the distinct regulatory mechanisms operating at each stage. This approach allows incorporation of transcriptional regulation by factors such as Fur and Fnr, as well as post-transcriptional regulation, including *RyhB*-mediated mRNA degradation, which occur on different timescales and are critical to the iron regulatory response in *E. coli*. With the exception of Fur, which is auto regulated and central to the iron regulatory network, we do not model separate regulatory pathways for proteins derived from the Semsey, et al. framework. Instead, transcription factors like Fnr and the cAMP receptor protein Crp, both significant in iron and oxidative stress regulation, are incorporated into our model (72,73). For the regulatory proteins described below (Equations 11–18), we explicitly model transcription and translation as separate steps. Messenger RNA and protein production are modeled using Michaelis–Menten kinetics, with production terms accounting for transcription factor regulation, dilution terms for each transcript, and additional degradation terms where applicable. In these equations, Fur-Fe²⁺ repression appears directly in the transcription term, and the ribosomal translation rates for siderophore associated proteins are scaled by ε_*τ*_. *K*_*d*,*FeFur*_^*ei*^ represents the dissociation factor of inhibitory *Fur-Fe^2+^* at the binding site of gene *e_i_*. Although *K*_*d*,*FeFur*_^*ei*^ should in principle vary among transcripts, in the implemented model we applied a uniform value *K*_*d*,*FeFur*_^*ei*^ = *K*_*f*_ throughout as a simplifying assumption, consistent with reported Fur binding affinities (68).

Transcription of *entF* and *feoB* is described in Equations 11 and 12, respectively.

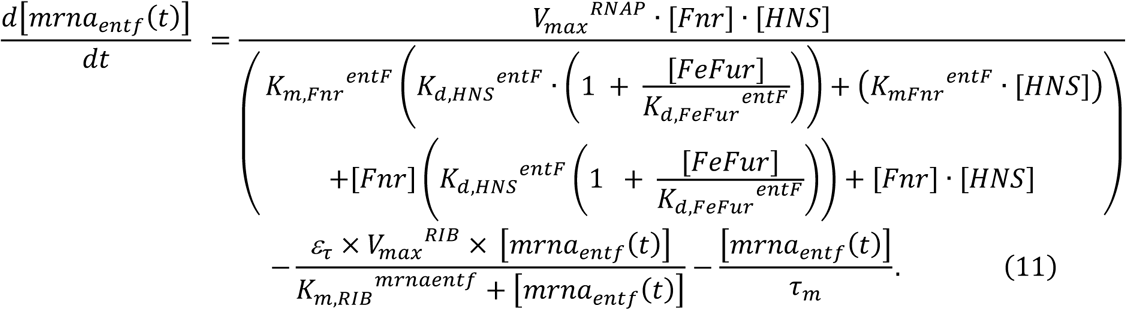

Equation 11 models the dynamics of *entF*. The first term represents *entF* transcription, where Fnr functions as a dual regulator to activate the fes-ybdZ-entF-fepE operon. Transcription is modulated by competition at the promoter between inhibitory Fur-Fe²⁺ and HNS; the parameters *K*_*d*,*HNS*_^*entF*^and *K*_*d*,*FeFur*_^*entF*^ denote their respective binding affinities, and *K*_*m*,*Fnr*_^*entF*^ is the Michaelis–Menten half-maximal constant for Fnr. The second term accounts for translation of the transcript into protein with ribosomal *V*_*max*_^*RIB*^scaled by ε_*τ*_. The final term represents mRNA degradation and dilution over one cell generation, represented by *τ*_*m*_ (11).

Equation 12 models the transcription dynamics of *feoB*. Fnr activates transcription for the *feoABC* operon, while Fur-Fe²⁺ inhibits it. In the production term, the effective transcription rate is modulated by Fnr relative to its binding affinity *K*_*m*,*Fnr*_^*feoB*^ and by Fur-Fe²⁺ relative to its binding affinity *K*_*d*,*FeFur*_^*feoB*^. The second term represents ribosome-mediated translation of mRNA, and the third term accounts for mRNA dilution during cell division *τ*_*m*_.

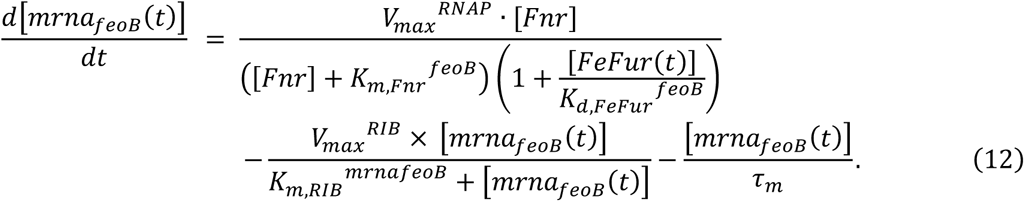

Transcription dynamics for *entB* are described in Equation 13:

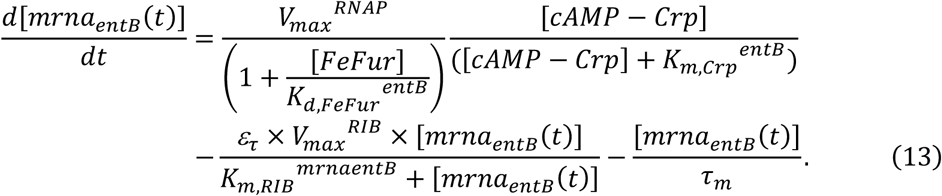

Here, cyclic AMP-activated Crp promotes transcription initiation for the entCEBA-ybdB operon, and we assume a sustained cAMP supply under our modeled conditions. *K*_*m*,*Crp*_^*entB*^ is the half-maximal constant for Crp-mediated activation of entB transcription, and Fur-Fe²⁺ inhibits transcription via its dissociation constant *K*_*d*,*FeFur*_^*entB*^. The second term captures translation of mRNA, and the third term accounts for mRNA dilution during cell division.

Equation 14 models *fepA* transcription dynamics.

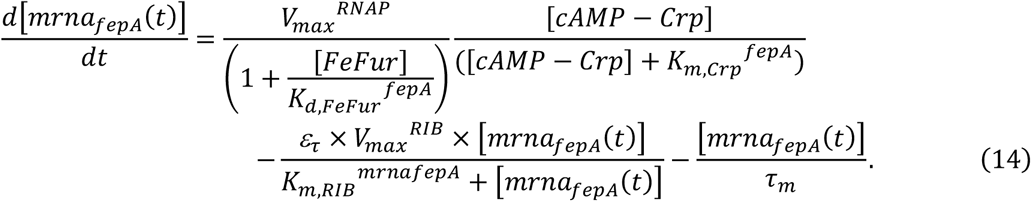

Here, *K*_*m*,*Crp*_^*fepA*^ is the half-maximal constant for *fepA* transcription. Under iron-sufficient conditions, active Fur-Fe²⁺ represses *fepA*, but when iron is limited, this repression is lifted, enhancing the uptake of iron-enterobactin complexes. Under these iron stress conditions, cells also upregulate esterases to hydrolyze and break down the complexes. The second and third terms account for translation and dilution of mRNA, respective.

Equation 15 models the dynamics of iron-enterobactin esterase (*fes*) transcription similar to the form used for *entF*.

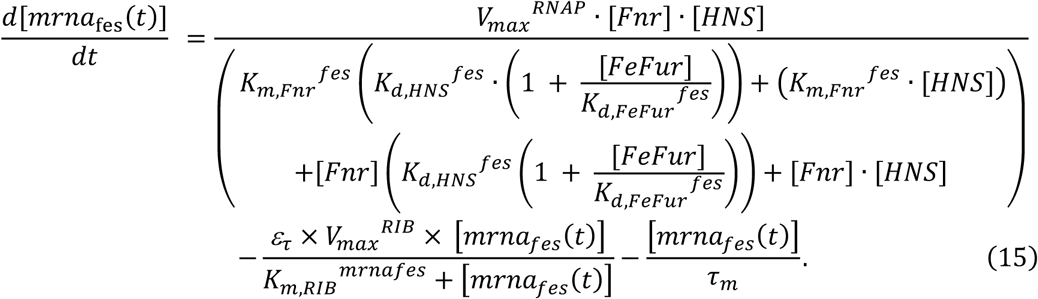

Here, transcription is activated by Fnr in conjunction with HNS, with the effective transcription rate modulated by the binding of Fur-Fe²⁺, with RNA polymerase activity modulated by regulatory parameters *K*_*m*,*Fnr*_^*fes*^, *K*_*d*,*HNS*_^*fes*^, and *K*_*d*,*FeFur*_^*fes*^. The last two terms capture protein synthesis from mRNA and dilution during cell division.

Equation 16 describes the dynamics of *sodA*, which encodes manganese superoxide dismutase, a key enzyme in the iron metabolic network. *SodA* is transcriptionally regulated by Fur (74) and is further targeted by *RyhB* for degradation. However, *SodB*, the other cytoplasmic superoxide dismutase that requires iron as a cofactor, is not regulated by iron levels. *E. coli* deploys these enzymes in a stress- and target-specific manner (75,76), with *SodA* generally providing more effective protection against DNA damaging radicals (77). Equation 16 models the dynamics of s*odA* mRNA production:

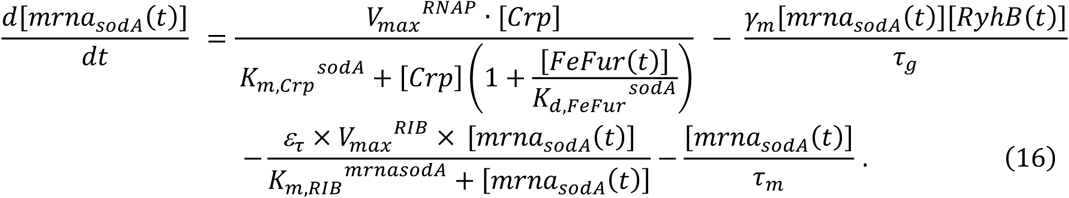

Equation 17 models *Fur* transcription as follows:

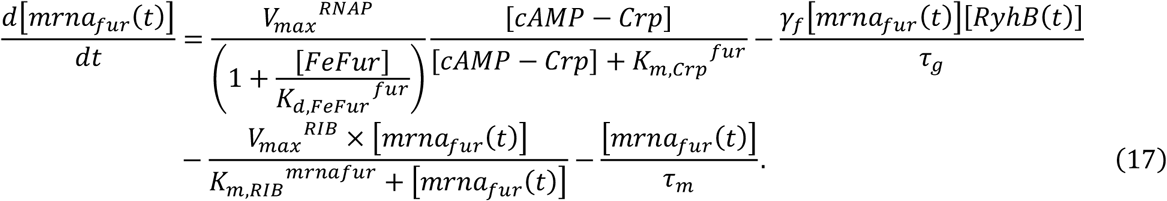

In this formulation, Fur auto-regulates its own expression by binding to its promoter in the presence of iron, while indirect regulation via *RyhB* reduces Fur transcription when iron is not limiting.

For each transcript in our model, denoted by m_B_ where n represents the gene (e.g., *entF, feoB, entB, fes, sodA,* or *fur*), protein production is described using a two-step process. The translation of each transcript is modeled by Equation 18:

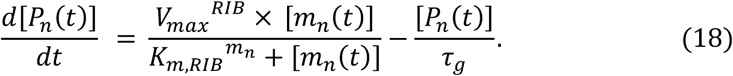

Michaelis–Menten kinetics are used to capture the enzymatic activity of the ribosome in translation. We model the dynamics of *RyhB*, *iscRSUA* transcripts, and the protein Suf following the approach established in the Semsey, et al. model.

#### 4.2.4 Extracellular Model of Redox Molecules and Volume Exchange

The extracellular environment model simulates iron cycling and redox balance, capturing the complex interplay between iron availability and redox dynamics. After synthesis, cells export siderophores into the extracellular space, contributing to the measured extracellular siderophore variable *ENTER*_*e*_(*t*).

This export is modeled in Equation 19.

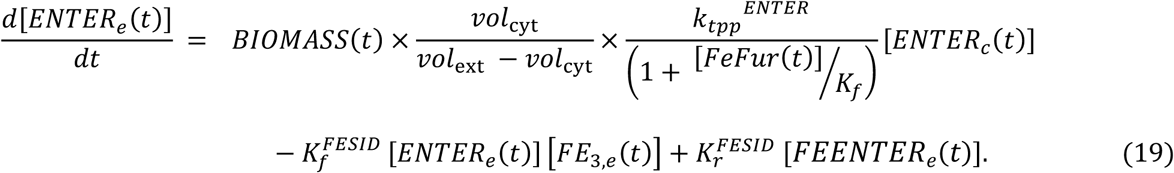

The first term represents a mass term for siderophore export that is scaled by cell density (biomass) and a volumetric compensation factor 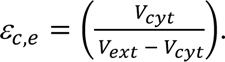 The second and third terms are mass action reactions describing the chelation of iron by siderophores and the subsequent dissociation respectively.

The binding constant *K_f_ ^FESID^* is adjusted to reflect competition from other siderophores and host iron chelators such as lactoferrin, transferrin, and ferritin; 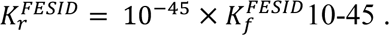

Extracellular iron dynamics are modeled separately for ferrous and ferric iron in equations 20 and 21. Reactions include recycling processes accounting for the oxidation of Fe²⁺ and reduction of Fe³⁺ by oxidants (e.g., hydrogen peroxide) and cellular reductants (modeled as a composite of NADPH and glutathione). Additionally, as cells die, the partially contribute iron to the extracellular nutrient pool, which is modeled by scaling the cell death term with a nutrient conversion factor *k*_*fr*_. The ratio of recycled ferric and ferrous iron is set by η (78).

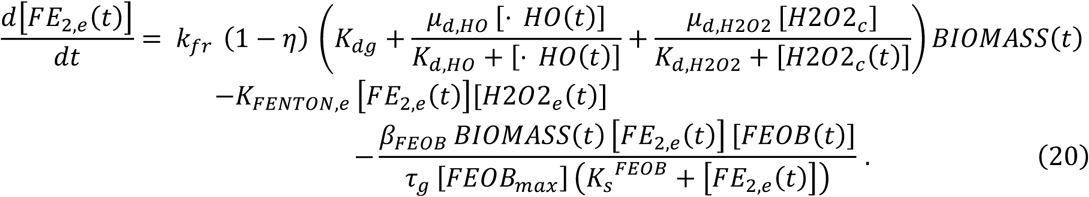

The first term in Equation 20 accounts for iron recycling from cell death, where *k*_*fr*_ scales the iron released, and the fraction (1 − *η*) designates the portion recovered as ferrous iron. The compound term is derived directly from the growth equation (see section 4.1.9) representing death rate in response to toxicity caused by oxidative stress. The second term represents extracellular Fenton chemistry consuming *Fe^2+^* via reaction with hydrogen peroxide. The third term captures ferrous iron uptake mediated by the *FeoB* permease, scaled by biomass and uptake capacity.

Extracellular ferric iron dynamics is modeled in Equation 21:

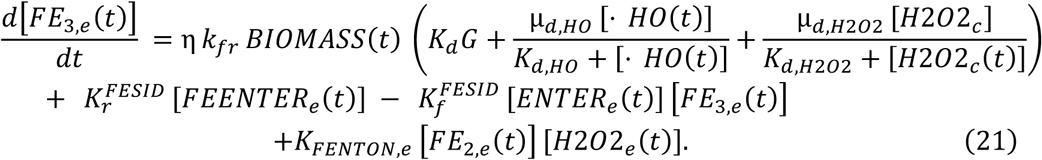

The first term captures ferric iron recycling from dead cells. The second and third terms describe the dissociation and binding kinetics of ferric iron with extracellular enterobactin. The last term accounts for ferric iron production through the extracellular Fenton reaction between Fe²⁺ and hydrogen peroxide.

Extracellular iron-enterobactin complexes re-enter the cytoplasm via the FepA uptake machinery, an ABC transporter whose rate is assumed to be limited by the available outer membrane protein FepA. The uptake velocity is scaled by cell density (BIOMASS) and FepA availability. Equation 22 describes extracellular ferric-enterobactin dynamics:

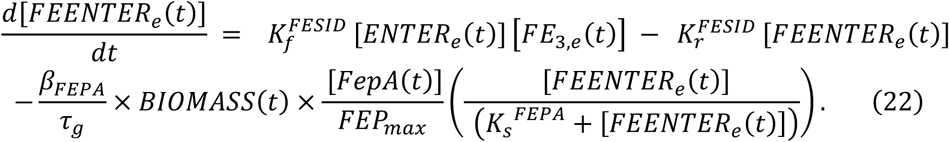

The first term represents complex formation through the binding of extracellular enterobactin (*ENTER*_*e*_) with ferric iron (*FE*_G,*e*_) using the binding constant *K_f_*^*FESID*^. The second term accounts for complex dissociation (rate constant *K_r_*^*FESID*^); and the third term models FepA-mediated transport scaled by cell density.

Equation 23 describes intracellular ferric enterobactin dynamics where the first term captures uptake from the extracellular compartment, which is adjusted by the extracellular to cytoplasmic volumetric compensation factor 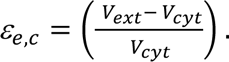 The second term represents the reduction of ferric-enterobactin by enterobactin esterase (with catalytic constant *K*_*kcat*_^*ENTERES*^, while the third term accounts for dilution due to cell division.

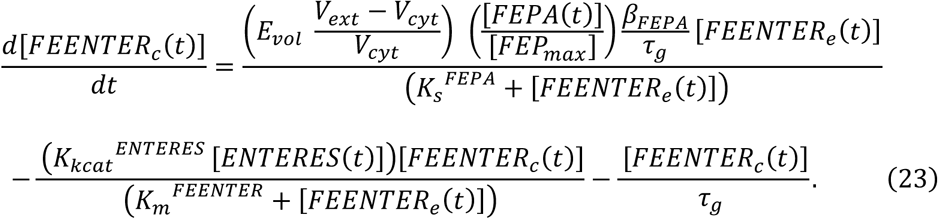

Notably, our β parameters *β*_*FEPA*_ and *β*_*FE*O*B*_ are based on extracellular iron concentrations and scaled by the cellular volumetric ratio, differing from the *β* parameter in the Semsey, et al. model.

#### 4.2.5 Peroxide Flux and Oxidative Stress Model

We captured the interplay of iron regulation and oxidative stress in *E. coli* by incorporating a flux model from Uhl et al. (8, 10). This model tracks both intra- and extracellular peroxide levels, simulates peroxide transport, and includes the superoxide dismutation pathways. Here, redox interactions, peroxide diffusion, and superoxide dismutation are modeled using mass action kinetics, while catalase- (KAT) and peroxidase (AHP)-catalyzed decomposition of hydrogen peroxide are represented via Michaelis–Menten kinetics. Our peroxide model also replicates periodic hydrogen peroxide injections following a kinetic protocol described in Supplementary Materials Appendix S6 (Figure S10). In our experimental multi-stress model (described in Section 4.1), 0uM or 100 μM hydrogen peroxide was periodically added to *E. coli* grown in M9 culture media with sufficient (approximately 7.2 μM) or depleted (approximately 0.072 μM) iron, with exogenous peroxide stress added every 90 minutes over a 12-hour period. In our computational model, a pulse function, ℎ_*e*_(A,t), replicates these discrete additions of extracellular peroxide, allowing simulation of bacteria response to four distinct conditions: sufficient iron (R), depleted iron (D), sufficient iron with peroxide pulses (RM), and depleted iron with peroxide pulses (DM). The pulse function is illustrated in supplementary materials (Figure S10).

#### 4.2.6 Metabolic Model of Hydrogen Peroxide Flux

Our model considers two sources of hydrogen peroxide: production via cellular metabolism and diffusion from the extracellular space. During cellular respiration, molecular oxygen (O_2_) is partially reduced to produce superoxide (O_2-_) and hydrogen peroxide (*H*_2_*O*_2_). Superoxide is subsequently converted to *H*_2_*O*_2_ via the action of superoxide dismutases (SPOD). Equation 24 describes intracellular H₂O₂ dynamics.

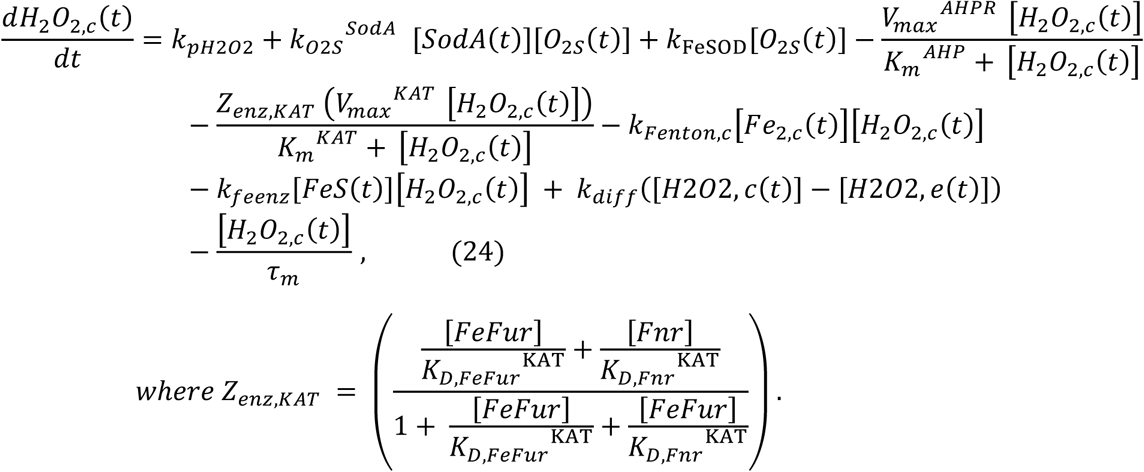

The first term (*k*_*pH*2O2_) represents spontaneous production. The second and third terms represent dismutation of superoxide by cytoplasmic dismutases *SodA* and *SodB* (FeSOD) respectively. The fourth and fifth terms capture *H*_2_*O*_2_ scavenging via catalase and peroxidase activities, modeled using Michaelis–Menten kinetics and modulated by a regulatory factor (Z_enz,KAT_ term) reflecting Fur and Fnr control. The sixth and seventh terms represent *H*_2_*O*_2_ consumption through Fenton reactions with free ferrous iron and with iron–sulfur clusters, respectively. The final terms reflect peroxide diffusion and dilution due to cell division.

Equation 25 models superoxide dynamics, with the first term representing production and the subsequent terms representing its dismutation and dilution.

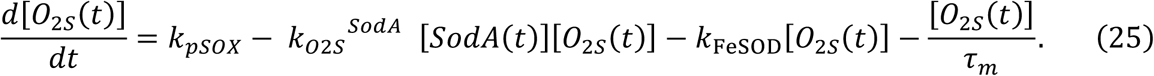

The dynamics of hydroxyl radical in our model is shown in Equation 26.

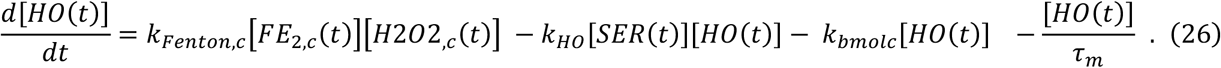

the first term reflects its production via Fenton chemistry between free ferrous iron and *H₂O₂*, the second term models its reaction with serine (9, 80), and the third term represents its overall oxidative attack on other cellular biomolecules (8,9). The final term reflects dilution due to cell division.

Extracellular *H₂O₂* dynamics are described by Equation 27.

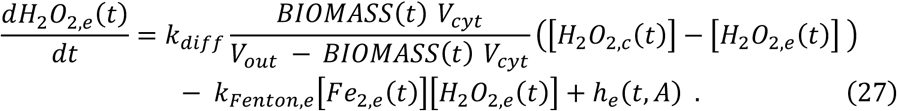

The first term represents diffusion of H₂O₂ from the cytoplasm to the extracellular space, scaled by cell density (BIOMASS) and the cytoplasmic-to-extracellular volume ratio. *V*_*cyt*_ represents the cell internal volume and *V*_*out*_ corresponds to the total volume and is set to 1 mL to correspond to experimentally determined cell density (CFU/ml) units. The second term captures H₂O₂ consumption in extracellular Fenton reactions, and the third term is a pulse function, ℎ_*e*_(*t*, *A*), that simulates repeated bolus injections of hydrogen peroxide at the intervals specified in the experimental stress response model. This hydrogen peroxide spiking model is shown in supplementary Figure S10.

#### 4.2.7 Intracellular Iron Model and Volume Exchange

We expanded the original Semsey, et al. iron model to distinguish intracellular ferrous iron that drives growth and ferric iron stores, which led to the use of two equations to describe iron flux. Equation 28 captures intracellular ferrous iron dynamics.

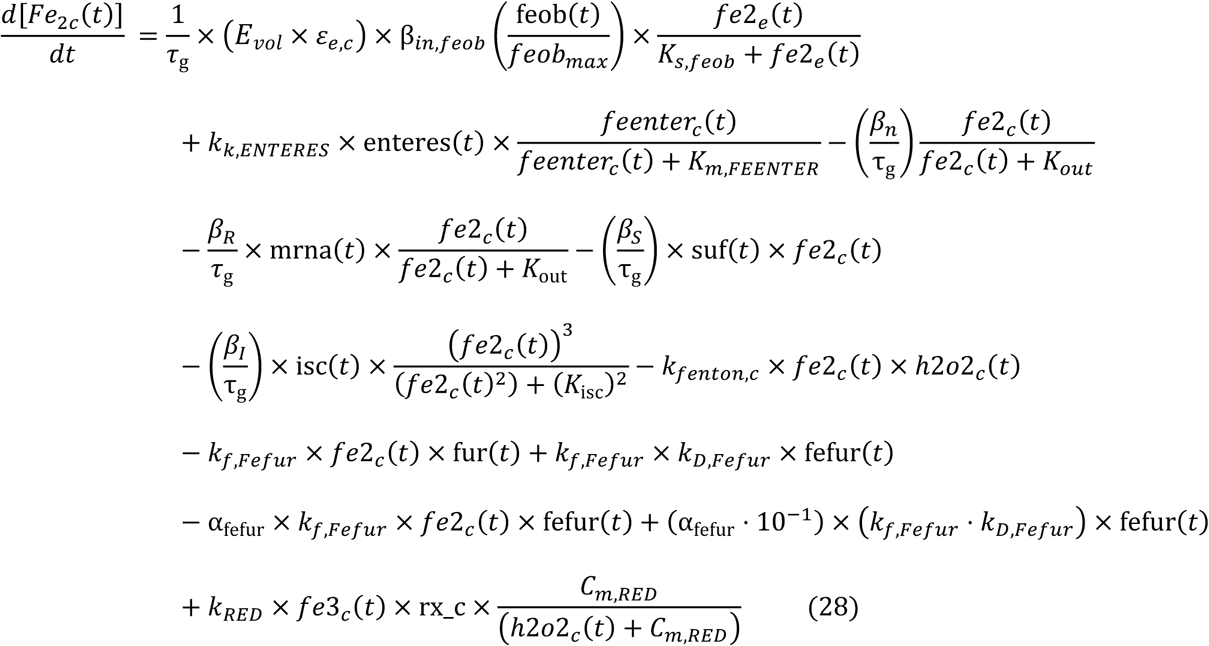

The first term represents ferrous iron uptake while the second term accounts for ferric iron uptake, thus expanding the original unified uptake mechanism used in the Semsey model. The third through sixth terms (from the Semsey model) describe processes related to iron utilization and distribution. The seventh term represents ferrous iron oxidation, and the eighth and ninth terms capture free iron exchange with the ferric uptake regulator. The final term models the recycling of ferric to ferrous iron via cellular esterases, an effect modulated by peroxide activity.

Equation 29 models intracellular ferric iron dynamics.

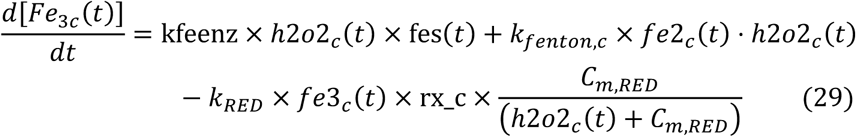

The first term represents the release of ferric iron due to peroxide-induced damage to iron-sulfur clusters. The second term accounts for ferric iron generation via the Fenton reaction between ferrous iron and hydrogen peroxide. Finally, the third term describes the recycling of ferric iron back to ferrous iron mediated by cellular esterases and modulated by peroxide activity.

#### 4.2.8 Bacterial Growth Model Linked to Redox and Intracellular Iron Stress

We developed a bacterial growth model that incorporates the effects of both iron and peroxide stress on cell proliferation and death. Our model uses a Monod-like formulation for cell density (BIOMASS, Equation 30) that combines the positive influence of intracellular ferrous iron with the inhibitory effects of oxidative stress. Peroxide-induced cell death is modeled via two mechanisms - direct peroxide toxicity and radical production through Fenton Chemistry (8,81).

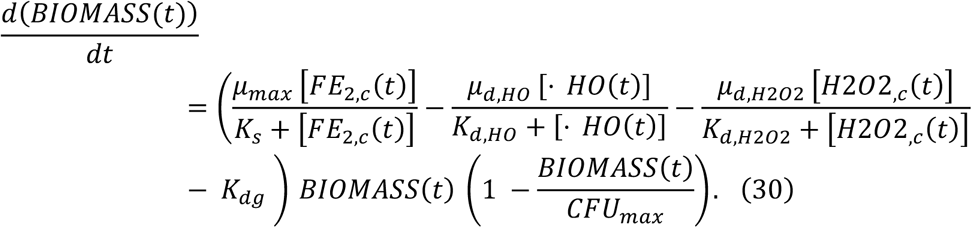

In Equation 30 the first term represents intracellular ferrous iron associated growth, while the second and third terms capture the inhibitory effects of hydroxyl radicals and hydrogen peroxide respectively. *K*_*d*g_ accounts for additional death processes not associated with redox modalities in our model. The term 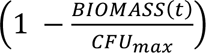 is a logistic growth constraint that enables predicted BIOMASS values to correspond to the range of experimentally observed CFUs over a 12-hour period for each condition (53).

Cell division time *τ*_*g*_ is modeled as a variable and is dynamically linked to cell density via Equation 31:

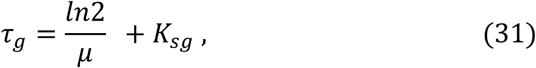

with the net growth rate *μ* defined as

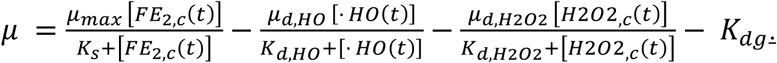

*K*_*s*g_ is the cellular maintenance coefficient, the growth rate required to maintain the cell population size, and is set to 0.02/min (81). Although a nominal doubling time of approximately 36 minutes (as used in the Semsey model) serves as a baseline for healthy cells, our model dynamically adjusts *τ*_*g*_ in response to varying iron and peroxide conditions.

The complete assembled model consisted of 50 coupled ordinary differential equations with 114 parameters (see supplementary files Supplementary Equations and Supplementary Spreadsheet – Model Specification). All computations were performed in MATLAB R2022b (MathWorks, Natick, MA) using the ODE15s solver with tolerance levels set to 1e-8.

### 4.3 Model Parameterization

We optimized our model parameters using empirical data from in vitro dual-stress experiments in *E. coli* described in Section 4.1 (19,53). Four phenotypes were examined: iron-replete (R), iron-deplete (D), iron-replete with 100 µM peroxide (RM), and iron-deplete with 100 µM peroxide (DM), and five response variables were measured under each condition – extracellular siderophore levels (enter.e), intracellular iron concentration (fe2.e), extracellular iron concentration (fe2.e), extracellular peroxide (h2o2.e) concentration, and cell density (biomass). To robustly parameterize the model for multi-phenotype predictions, we employed the multi-phenotype parameterization workflow shown in Figure 2. First, we performed sensitivity analysis using a composite set of error metrics to identify critical system drivers across diverse phenotypes and classify them into significance class (I to III) (19, 66). Next, we sequentially optimized identified parameters based on their significance classes. Finally, we used ensemble modeling to generate aggregate models that effectively capture iron regulation under multiple stress phenotypes.

#### 4.3.1 Augmented Sensitivity Analysis

In previous work (19), we introduced a sensitivity analysis approach that used an augmented set of error measures to robustly identify key drivers in the *E. coli* iron regulatory network. Instead of relying solely on average response values, our method incorporated additional error measures – including average absolute gradients (first derivatives), average absolute Laplacians (second derivatives), and maximum absolute gradients—to capture dynamic behaviors across each simulated stress phenotype. In this work we used the same approach (Figure 14) and generated 45,200 parameter sets using Latin Hypercube Sampling (LHS) and computed partial regression correlation coefficients (PRCCs) using DAKOTA (82) (Sandia National Laboratories). A t-statistic corresponding to a p-value of 0.001 was used as the significance threshold (83). We performed the analysis under our four simulation conditions, replete, replete with hydrogen peroxide stress, deplete, and deplete with hydrogen peroxide stress. Simulation results were used to identify the parameters that most significantly influence model variables across the multi-phenotype landscape, which were then designated as critical for optimization.

**Figure 14:**
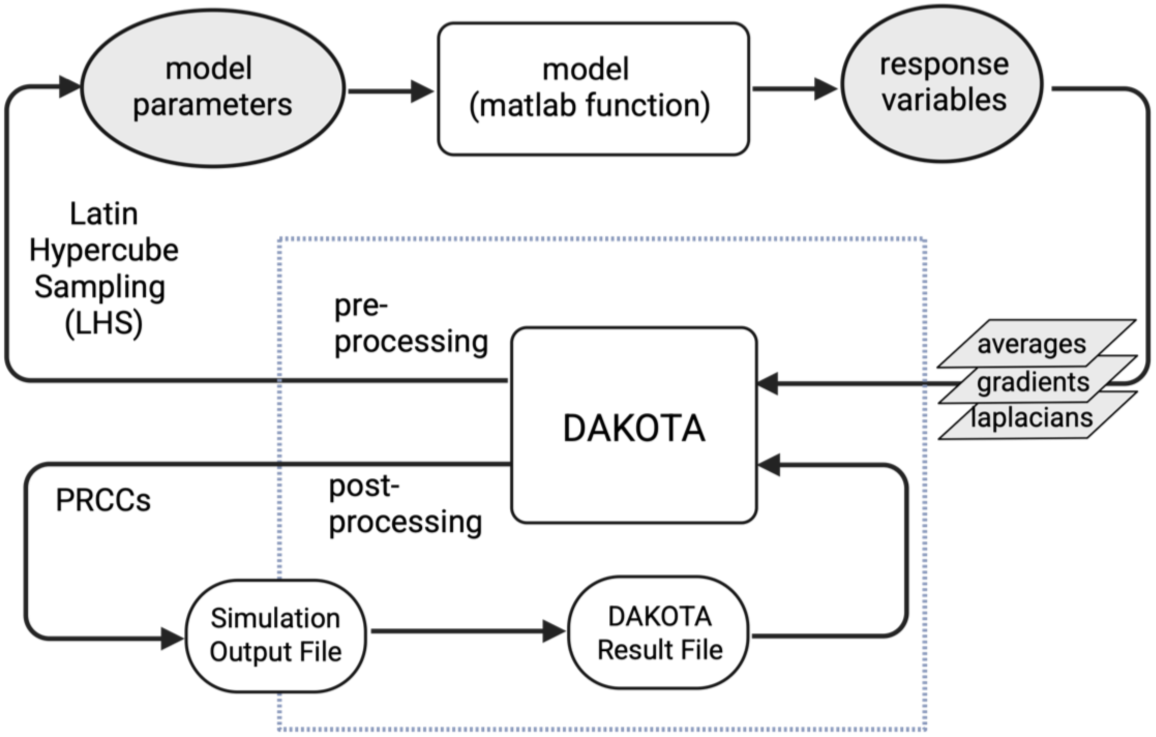
Augmented Measures Sensitivity Analysis. Schematic of the computational pipeline for global sensitivity analysis. Model parameters sampled via Latin Hypercube Sampling (LHS) are passed to the MATLAB model to generate response variables. Augmented error measures (averages, gradients, and Laplacians) are computed and processed through DAKOTA to yield PRCCs that identify influential parameters.

#### 4.3.2 Multi-Phenotype Optimization

The optimization objective for this study was to identify a parameter set that consistently fits our four stress phenotypes (R, D, RM, DM) and five response variables (enter.e, fe2.e, fe2.e, h2o2.e, biomass). We conducted a computational experiment comparing three optimization strategies: multi-phenotype multi-response (MPMR), which integrated data across all stress phenotypes and response variables; single-phenotype multi-response (SPMR), which utilized multiple response variables from one phenotype; and single-phenotype single-response (SPSR), which focused on a single response variable within a single phenotype. Datasets from three of four stress conditions (R, D, RM) were used for optimization and validation, while the iron-deplete with peroxide stress condition (DM) was held out for testing the optimized model. Because DM combined iron limitation with oxidative stress, it represented a true dual-stress condition, posed a more severe biological challenge to bacterial viability, and was expected to be difficult to predict computationally compared to single-stress environments. Response data was further categorized into three groups: metabolic (intracellular iron and extracellular siderophore levels), environmental (extracellular iron and hydrogen peroxide levels), and phenotypic (cell density via biomass). Candidate models (parameter sets) were evaluated using an augmented set of error measures, including sum-of-squares, gradient, and Laplacian errors, to determine which optimization strategy best captured the entire set of multi-phenotype responses. Appendix S7 provides a summary of the data combinations used for the multi-phenotype optimization analysis.

#### 4.3.3 Practical Identifiability and Sequential Estimation

Relying solely on initial estimates derived from incomplete biochemical data is insufficient to establish reliable parameter bounds. Instead we implemented a stepwise parameter estimation method that iteratively explored the parameter space (θ), to identify practical bounds for each parameter based on the training sets (45,84). Because this approach was computationally intensive, we increased the sampler’s step size to improve efficiency. We coupled this approach with sequential optimization (36) where parameters were optimized sequentially based on their sensitivity to model variables. The optimal value and distribution determined from a prior sequence of optimization (of a class of more influential parameters) was used as the initial set for subsequent optimization steps. Each estimation was carried out over 14 generations using DAKOTA’s JEGA-based multi-objective genetic algorithm (MOGA) for different combinations of responses for each optimization strategy (MPMR, SPMR, or SPSR). To standardize the analysis across all result sets from each optimization strategy, the minimum number of solutions across the Pareto ranks was adopted as the solution size for each group. To evaluate the errors associated with each model regardless of the phenotype and response group it was drawn from, we recomputed a normalized multi-phenotype error value which computed the deviation from ground truth temporal data and coverage value which measured a model’s predictive value based on significance testing (for a given response, a non-temporal comparison of all measured and predicted values). We computed the rank of each model by assigning a composite score based on its error and coverage value. The post-optimization processing (4,000 – 8,000 simulations per optimization strategy) and ranking was implemented using MATLAB on an Apple MacBook Pro (32GB, 10-core Apple M1 Pro chip) while the sequential estimation pipeline (200,000 – 540,000 evaluations per optimization strategy) was executed on a highly parallel 48-core Carya cluster computing system (189GB, Intel Xeon G6252 CPU; Research Computing Data Core at the University of Houston). Appendix S8 provides a summary of the approach we used for identifiability and sequential estimation.

#### 4.3.4 Ensemble Modeling

We applied ensemble modeling to identify combinations of models within each optimization strategy that approximated observed data under varying stress conditions and inputs. The ensemble selection process was formulated as a linear programming problem (Table S2, Appendix S9) that selects an optimal subset of models that minimized error along the ensemble dimension; subsets were selected from the population of candidate models generated by each optimization strategy. Additionally, error weighting was applied as a technique to differentially weight error values of response components, allowing the biasing of ensembles toward specific predictive objectives. For each ensemble, we computed an aggregate error score using weighted averaging, where models with lower errors contributed more significantly to the predicted model outcome and the influence of models with higher errors was reduced. Our ensemble modeling with differential error weighting approach is generalizable and can be used to prioritize accuracy in predicting specific response variables or phenotype outcomes versus overall accuracy. Combined with MPMR optimization, generating tunable ensembles to application or phenotype specific use cases becomes highly feasible.

## Acknowledgments

The authors would like to thank Jeffery Sarlo and other members of the Research Computing Data Core at the University of Houston for technical assistance and support provided through the Carya computing cluster. We would like to thank former members of the May lab at the University of Houston for technical assistance generating experimental data used in this study. Research supported in part by National Science Foundation grants MCB1445470, MCB1812455, PHY2019745, MCB2227598, and National Institute of Health grant P41EB023912.

